# RIG-I-like receptor-dependent type I Interferon regulates antigen dose and activation in yellow fever vaccine 17D-infected antigen presenting cells

**DOI:** 10.1101/2025.10.08.680190

**Authors:** Magdalena Zaucha, Elena Winheim, Antonio Santos-Peral, Apurva Dhavale, Paul Schwarzlmueller, Frank Dahlstroem, Giulia Spielmann, Magdalena K. Scheck, Linus Rinke, Yiqi Huang, Varvara Arzhakova, Katharina Eisenächer, Hadi Karimzadeh, Michael Pritsch, Julia Spanier, Ulrich Kalinke, Julia Thorn-Seshold, Giovanna Barba-Spaeth, Simon Rothenfusser, Anne B. Krug

## Abstract

The live-attenuated yellow fever vaccine 17D-204 (YF17D) activates robust innate immune responses followed by rapid induction of adaptive immunity resulting in long-lasting protection. YF17D triggers the production of type I interferons (IFNs) which have a dual role in antigen presenting cells regulating their infection and contributing to their activation. Infection with YF17D was detected in primary human blood monocytes and conventional dendritic cells (DCs) and in monocyte-derived DCs but was highly restricted by type I IFN. Blocking IFNAR signaling in YF17D-infected PBMC from vaccinated donors resulted in increased activation of YF17D-specific CD8^+^ T cells. Consistently, peak IFN-alpha plasma levels correlated inversely with the CD8^+^ T cells response in YF17D vaccinees. Loss of function experiments demonstrated a dominant role of retinoic acid inducible gene I (RIG-I)-like receptors (RLRs) and mitochondrial antiviral signaling protein (MAVS) for type I IFN induction and restriction of YF17D. The type I IFN response was mediated by 5’ tri- or diphosphate dsRNA intermediates that are formed during YF17D infection. *In vivo* proximity labelling (IPL) of RIG-I and next-generation sequencing confirmed interaction of RIG-I with YF17D-dsRNA in infected cells. Thus, YF17D-triggered RLR-signaling restricts viral replication through type I IFN and thus limits the production of viral antigens that can be presented to T cells.

## Introduction

The yellow fever vaccine 17D-204 (YF17D) induces long lasting protective humoral and cellular immunity against infection after a single vaccine dose and thus is one of the most effective vaccines available. Administered to over 850 million people globally (1), it is considered the gold standard of an effective vaccination. The live-attenuated YF17D vaccination leads to productive but limited infection in humans as indicated by a transient viremia which peaks on day 5 to 7 after vaccination (2, 3). The viremic phase is accompanied by a robust induction of type I IFN stimulated genes in peripheral blood mononuclear cells (PBMC) indicating a temporary systemic innate antiviral response (4, 5). The initial viral load was shown to determine the magnitude of the subsequent CD8^+^ T cell response (2), implying the importance of a sufficient amount of viral antigen to induce adaptive immunity. At the same time restriction of YF17D replication by type I IFN is critical for the safety of this live-attenuated vaccine (6, 7). Type I IFN is also known to promote dendritic cell (DC) maturation and antigen presentation to T cells (8–14) and a robust type I IFN response is considered to be a hallmark of an effective vaccine (5). Indeed, YF17D vaccination induces a coordinated temporary response of circulating DCs and monocytes 3 to 7 days after vaccination dominated by upregulation of type I IFN induced genes. This includes genes involved in antigen processing and presentation as well as costimulatory molecules suggesting that type I IFNs elicited by YF17D promote the induction of adaptive immunity (15, 16). However, it was shown in mice that type I IFN can also reduce the amount of available antigen for presentation to T cells early after viral infection thereby dampening the T cell response (17). Thus, type I IFN induction in human antigen presenting cells (APCs) by a live viral vaccine such as YF17D may have opposing effects on the T cell response by promoting antigen presenting cell (APC) activation and by restricting viral replication thereby regulating the antigen dose.

Several studies showed productive but limited infection of human monocyte-derived DCs (mo-DCs) by YF17D inducing moderate upregulation of costimulatory molecules and production of type I IFNs and proinflammatory cytokines in vitro (18–22). Plasmacytoid DCs (pDCs) were shown to be primarily activated to produce type I IFN by immature YF17D particles or RNA transferred from other infected cells via Toll-like receptor (TLR) 7 and independent of viral replication, although direct infection of pDCs and induction of type I IFN via the cytosolic RNA sensor retinoic acid inducible gene (RIG)-I has also been described (23, 24). Sensing of YF17D by RIG-I-like receptors (RLRs) was also shown in human cell lines (5, 25–27). In mice, the activation of splenic DCs which are not productively infected by YF17D was shown to involve TLR2, TLR7 and TLR9 (22). It remains unanswered which viral sensors and downstream signaling pathways are responsible for the response to YF17D in human primary APCs besides pDCs and which pathways are relevant for controlling viral replication and production of viral antigens for presentation to T cells. In this study we investigated the interaction of human and murine APCs with YF17D and characterized the viral sensing pathways required for triggering type I IFN and restricting YF17D. We found that YF17D is restricted by RLR-dependent sensing of dsRNA intermediates leading to robust induction of type I IFNs which contribute to APC activation but limit availability of viral antigen for presentation to T cells.

## Results

### YF 17D infects human primary DCs and monocytes and is highly restricted by type I IFN

To compare the extent of YF17D infection between different primary cell types, PBMCs were isolated from healthy donors and infected with YF17D at 1 and 10 MOI for 48 hrs. Cell viability was not compromised up to this time point (Fig. S1). The percentage of cells that were labeled positive for the YF17D envelope (E) protein within different cell types was determined by flow cytometry (gating strategy shown in Fig. S2). Consistently, a dose-dependent increase in the percentage of E protein positive cells was observed in monocytes (HLADR^+^ CD88^+^ CD14^+^ and/or CD16^+^) and in conventional DCs (HLADR^+^ CD88^−^ CD14^−^ CD16^−^ CD11c^+^), while lymphocytes (T, B and NK cells) and pDCs did not show an E protein signal above background. Dose-dependent infection with YF17D was also observed in mo-DCs confirming previously published data and similar results were obtained using a Venus-YF17D reporter virus (Fig. 1B, gating strategy shown in Fig. S2A). Overlap of Venus fluorescence and E protein labeling was confirmed in infected Vero cells (Fig. S2B). Even at 10 MOI the infection rate in PBMCs (monocytes and cDCs) and in mo-DCs was limited suggesting viral restriction. While robust IFN-β induction was observed after infection with 1 MOI of YF17D, the same dose of heat-inactivated (HI) virus failed to induce IFN-β secretion in mo-DCs and monocytes (Fig. 1C). The addition of neutralizing antibodies against IFN-α, IFN-β and IFN-α/β receptor (IFNAR) significantly increased the percentage of Venus-YF17D^+^ mo-DCs and monocytes after infection at 1 MOI respectively (Fig. 1D and E) indicating that YF17D is highly restricted by IFNAR-signaling. Thus, YF17D efficiently triggers type I IFN production in monocytes and mo-DCs thereby limiting viral replication in these cell types.

**Figure 1.**
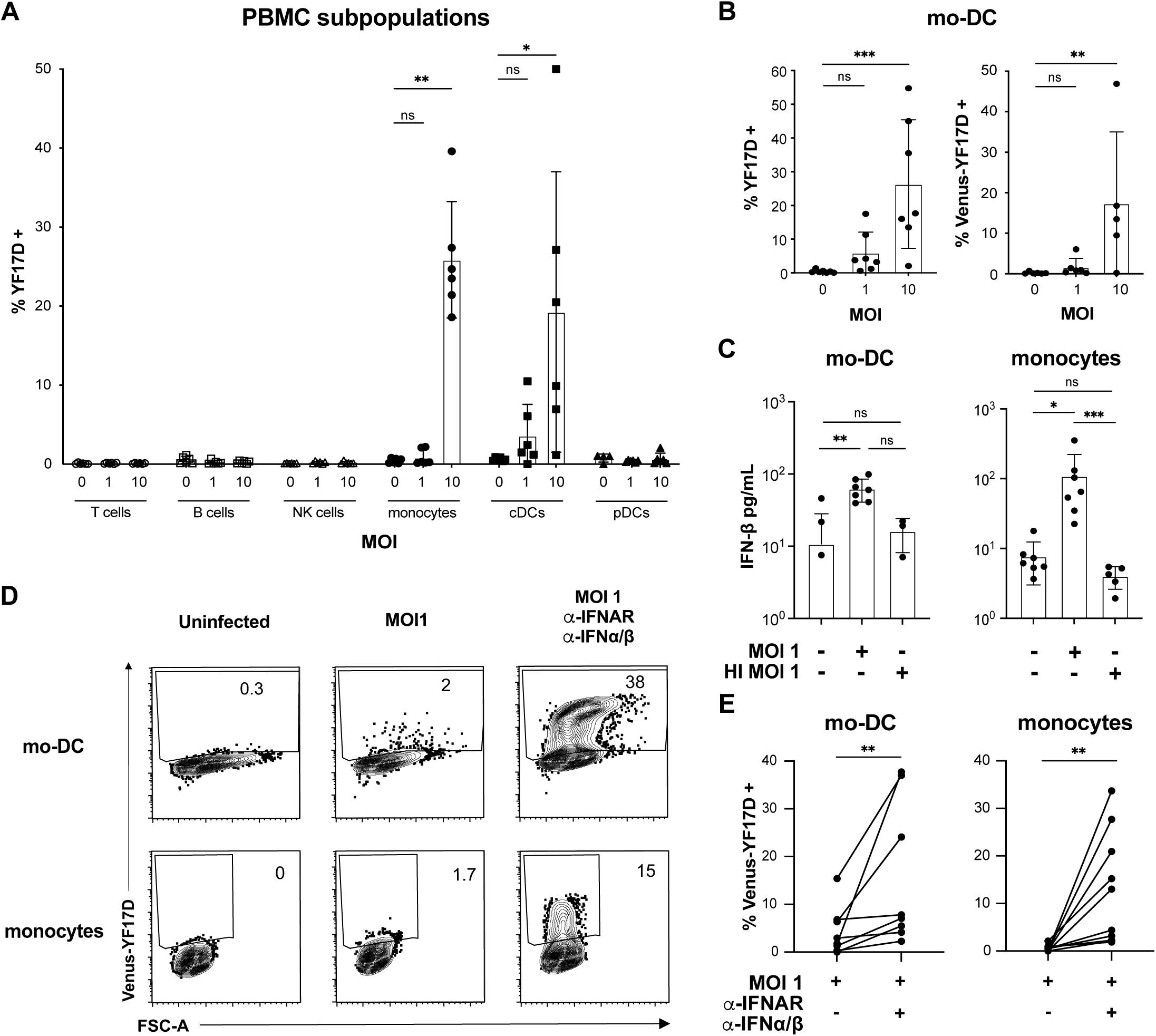
Infection of primary human DCs and monocytes with YF17D is restricted by type I IFN. (A) PBMC were infected with YF17D (MOI 1 and 10) or not infected. After 48 hrs of culture cells were analyzed by flow cytometry. Subpopulations were identified by surface marker staining and infected cells were identified by intracellular staining for the viral E protein using the 4G2 antibody. The percentages of E protein positive cells within the indicated subpopulations are shown as mean +/- SD (n=6) with individual data point indicated by symbols (*p<0.05, **p<0.01, ***p<0.001, Kruskal-Wallis-test with Dunn’s correction). (B) Mo-DCs were cultured with medium (control) infected at 1 or 10 MOI with YF17D (left panel, n=7, cells from 7 individual donors) or with Venus-YF17D for 36-48 hrs (right panel, n=4-5, cells from 5 individual donors). The percentages of E protein positive and Venus-positive cells are shown respectively (mean +/- SD; *p< 0.05, **p<0.01, ***p< 0.001, Kruskal-Wallis-test with Dunn’s correction). (C) IFN-β concentration in the supernatants of mo-DCs (n=3-7) and monocytes (n=5-7) infected with Venus-YF17D (MOI 1) or incubated with equivalent dose of heat-inactivated Venus-YF17D (HI) for 36 and 48 hrs respectively are shown (mean +/- SD; *p< 0.05, **p<0.01, ***p< 0.001, Kruskal-Wallis-test with Dunn’s correction). (D) Representative contour plots showing the Venus fluorescence signal after infection of mo-DCs and monocytes with Venus-YF17D (MOI 1) in the absence or presence of IFNAR blocking antibodies anti-IFN-α/-β neutralizing antibodies. (E) Percentages of infected mo-DCs (n=8) and monocytes (n=10) after infection with Venus-YF17D (MOI1) for 36 and 48 hrs respectively, in the absence or presence of IFNAR blocking antibodies and IFN-α/-β neutralizing antibodies are shown (mean +/- SD; **p<0.01, Wilcoxon paired test).

### YF17D infection induces activation of infected and bystander mo-DCs and monocytes

Type I IFN may contribute to the activation of APCs in an autocrine or paracrine manner. Exposure to live YF17D induced dose-dependent upregulation of activation markers such as PD-L1, CD80, CD86, and HLADR in mo-DCs and monocytes, whereas the corresponding induction levels after stimulation with LPS or R848 were even higher (Fig. 2A, B). Specifically, CD83 was moderately upregulated in mo-DCs (not significant, Fig. 2A), and CD86 was upregulated in both Venus-YF17D^+^ and Venus-YF17D^−^ mo-DCs with a trend towards a higher percentage of CD86^+^ cells in the Venus-YF17D^+^ cells (Fig. 2 C and D, not significant). The upregulation of activation markers in mo-DCs and monocytes in response to YF17D was slightly reduced but not abrogated by the addition of blocking antibodies against IFNAR and type I IFNs (Fig. 2 F and G, statistically significant for PD-L1 and HLADR in infected monocytes and for HLADR in infected mo-DCs). Thus, YF17D induces both direct and bystander activation in mo-DCs and monocytes, which is partially mediated by type I IFNs.

**Figure 2.**
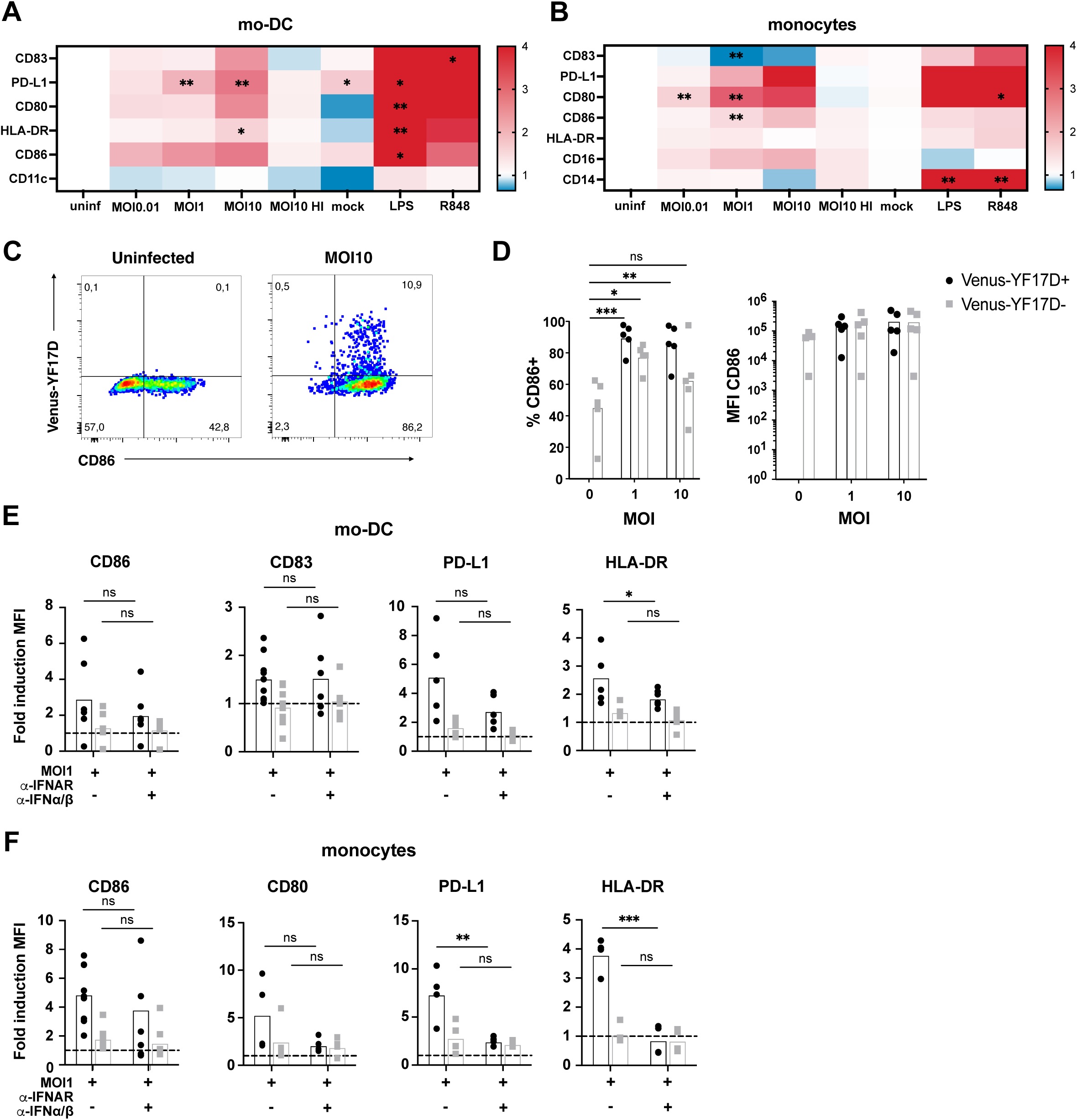
Activation of infected and bystander Mo-DCs and monocytes by YF17D. (A, B) Isolated mo-DCs (A) and monocytes (B) were infected with of Venus-YF17D (MOI 0.01, 1 and 10), heat-inactivated Venus-YF17D (MOI 10 HI) or incubated with a mock control obtained from sucrose gradient centrifugation for 36 and 48 hrs respectively. LPS and R848 were used as positive controls. MFI values of the indicated activation markers were normalized as fold-changes to uninfected/untreated controls. The color scale shows the mean fold-change values (n=3). White: unchanged, blue: downregulation, red: upregulation. Asterisks indicate significant p values (*p< 0.05, **P<0.01, Kruskal-Wallis-test with Dunn’s correction). (C) Exemplary dot plots showing CD86 expression and Venus-YF17D signal in mo-DCs infected with 10 MOI Venus-YF17D for 36 hrs or cultured in medium without virus. (D) Mo-DCs were infected with 1 or 10 MOI Venus-YF17D or cultured in medium for 36 hrs (uninfected condition). The percentages of CD86^++^ cells and the CD86 MFI values are shown for Venus^+^ (black) and Venus^−^ cells (grey) in the infected condition and for all cells in the uninfected condition (grey) as mean +/- SD with individual data points shown as symbols (n=5). Asterisks indicate significant differences to the uninfected control (*p<0.05, **p<0.01, ***p<0.001, One-way Anova with Dunnett’s correction). (E, F) mo-DCs (E) and monocytes (F) were infected with 1 MOI Venus-YF17D in the presence of absence of anti-IFNAR and anti-IFN-α/-β antibodies. Cells were analyzed for expression of the indicated activation markers by flow cytometry after 36 and 48 hrs respectively. MFI values for Venus-YF17D^+^ and Venus-YF17D^−^ cells were normalized to MFI values of uninfected cells (fold induction). Columns indicate the mean fold induction and symbols indicate individual data points (black: Venus-YF17D^+^; grey: Venus-YF17D^−^, n=6-10). Asterisks indicate significant differences between with and without IFN I blockade for Venus^+^ cells and for Venus^−^ cells respectively (*p<0.05, **p<0.01, ***p<0.001; 2-way Anova with Šidák‘s correction).

### Type I IFN limits CD8^+^ T cell response to YF17D

While type I IFNs contribute to APC activation their antiviral effect restricts YF17D limiting the availability of endogenous viral antigens for presentation to T cells. To study the net effect of YF17D-induced type I IFN on vaccine-induced T cell responses, PBMCs from 5 vaccinated donors and 2 unvaccinated donors were re-stimulated *ex vivo* with YF17D in the presence or absence of IFNAR blocking antibodies. As expected, IFNAR blockade resulted in enhanced YF17D infection of cDC and monocytes present in the donor’s PBMC (Fig. 3A, gating strategy shown in Fig. S3). CD86 and HLADR expression was upregulated in cDCs and monocytes in both conditions, but IFNAR blockade led to significantly lower expression of CD86 and HLADR in monocytes and CD86 in cDCs (Fig. 3B).

**Figure 3.**
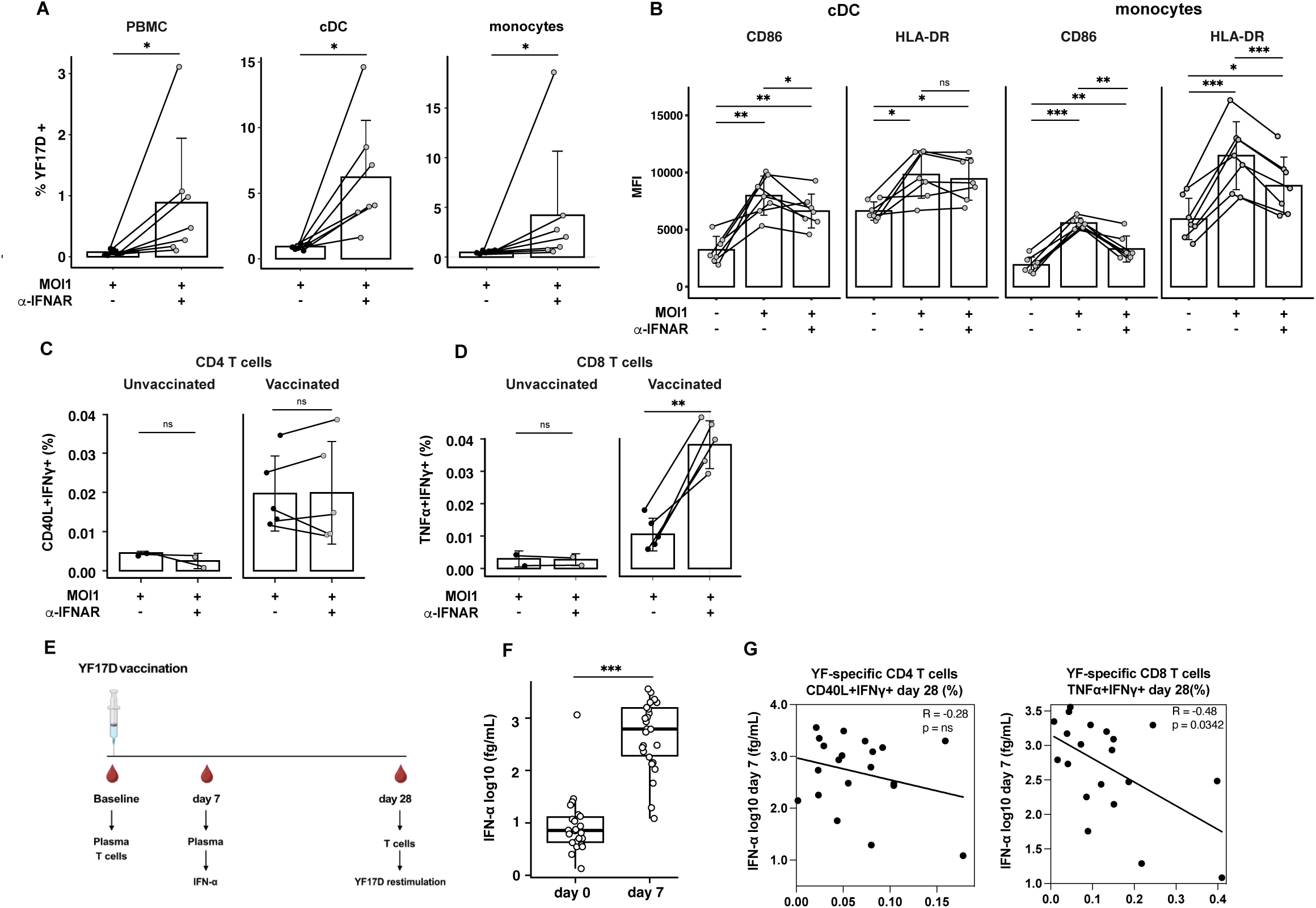
Type I IFN reduces YF17D-specific CD8^+^ T cell activation. (A-D) PBMC from healthy YF17D-vaccinated (n=5) and unvaccinated donors (n=2) were infected with live Venus-YF17D virus (MOI 1) in the presence or absence of 2.5 µg/mL of an anti-IFNAR blocking antibody with addition of Brefeldin A at 24 hrs followed by an incubation time of 20 hrs. (A) Percentage of Venus-YF17D positive cells of total PBMC, cDCs (gated as HLADR^+^ CD14^−^ CD11c^+^) and monocytes (gated as HLADR^+^ CD14^+^) shown as mean and SD (n=7). Asterisks indicate statistically significant differences (*p<0.05, paired Wilcoxon Matched pairs test). (B) MFI values of the indicated activation markers in gated cDCs and monocytes (mean +/- SD, n=7). Asterisks indicate statistically significant differences compared to the unstimulated control (*P<0.05, **P<0.01, ***P<0.001, paired t-test with Holm’s correction). (C-D) Frequency of YF17D-specific CD40L^+^IFNγ^+^ CD4^+^ T cells. (D) Frequency of YF17D-specific IFNγ^+^TNFα^+^ CD8^+^ T cells (mean +/- SD, n=2 for unvaccinated, n=5 for vaccinated). Asterisks indicate statistically significant differences (*p<0.05, **p<0.01, ***p<0.001, Wilcoxon matched pairs test). (E) Schematic representation of longitudinal PBMC and plasma collection of healthy vaccinees at baseline before and on days 7 and 28 after vaccination with a single dose of YF17D. Plasma samples were tested for cytokine concentration at baseline and on day 7. PBMCs from these donors were analyzed for the frequencies of YF17D-responsive CD4^+^ and CD8^+^ T cells on day 28. (F) Concentration of IFN-α2 in plasma samples (mean +/- SD, n=26, ***p<0.001, Wilcoxon matched pairs test). (G) Spearman rank correlation analysis between IFN-α2 plasma levels on day 7 and the frequencies of YF17D-responsive CD40L^+^IFNγ^+^ CD4^+^ T cells (left panel) and IFNγ^+^ TNFα^+^ CD8^+^ T cells (right panel) on day 28 (n=20).

The antigen-specific CD4^+^ and CD8^+^ T cell response was quantified by intracellular cytokine and CD40L staining (ICS) which allows the identification and quantification of antigen-specific CD4^+^ and CD8^+^ T cell responses after YF17D infection of PBMC from YF17D-vaccinated donors (28). The frequencies of IFNγ^+^TNFα^+^ CD8^+^ T cells in PBMC from vaccinees were increased when YF17D infection was performed in the presence of IFNAR blockade. This effect was not observed for antigen-specific CD40L^+^IFNγ^+^ CD4^+^ T cells (Fig. 3C, D and Fig. S3).

To further assess the impact of type I IFN responses on the T cell response *in vivo*, IFN-α levels were measured in plasma samples from 26 YF17D vaccinees before (day 0) and on day 7 after vaccination using an ultrasensitive immunoassay and YF17D-responsive T cells were quantified on day 28 (Fig. 3E). IFN-α plasma levels were significantly upregulated on day 7 compared to baseline (Fig. 3F) and IFN-α concentration on day 7 negatively correlated with the frequency of YF17D-specific TNF-α^+^ IFN-γ^+^ CD8^+^ T cells on day 28 after the vaccination (Fig. 3G). However, no significant correlation was observed with the frequency of YF17D specific CD40L^+^ IFN-γ^+^ CD4^+^ T cells induced by the vaccine. Thus, type I IFN limits T cell activation in response to YF17D, with a more pronounced impact on the CD8^+^ T cell response. This may be due to the restriction of virus propagation by type I IFN thereby reducing the availability of viral antigens for presentation to T cells.

### YF17D restriction by type I IFN depends largely on RLR/MAVS signaling

We then asked the question which receptors and pathways sense and restrict YF17D in APCs. Previous studies have demonstrated the involvement of multiple TLRs in murine DCs and the RLR/MAVS pathway in human pDCs and non-immune cells (22, 23, 25–27). However, the individual and combined contribution of TLR and RLR signaling to IFN-mediated YF17D restriction and the potential role of the cGAS/STING pathway have not been investigated so far. To address this question, we generated macrophages and DCs from bone marrow (BM) cells by culture with M-CSF (BM-DM) or GM-CSF cells (GM-DC/Mac) using BM cells of mice deficient in one or several signaling adaptors or IFNAR and infected them with 1 MOI of Venus-YF17D. By fluorescence microscopy, very few infected cells were detected in wildtype (WT) and MDA5 knockout (KO) BM-DM, whereas MAVS KO and IFNAR KO cells showed a higher frequency of infected cells (Fig. 4A). Flow cytometric analysis of CD11c^+^ MHCII^+^ cells from GM-CSF BM cultures allowed us to simultaneously examine two subpopulations, CD11b^hi^ MHCII^int^ GM-Mac and CD11b^int^ MHCII^hi^ GM-DC, as earlier described (29). Infection rates in GM-DC were 2 to 4 times lower than in GM-Mac, even when gating on these cell types within the same culture (Fig. 4 B and C). These results suggest that GM-DCs have a higher intrinsic resistance to infection with YF17D than GM-Mac. The percentage of Venus^+^ infected cells was significantly increased in IFNAR KO GM-Mac and GM-DC (p < 0.0001) compared to the wildtype (WT) control. Additionally, MAVS KO GM-Mac and GM-DC showed an increased percentage of infected cells (statistically significant for GM-DC, p = 0.0285), whereas MyD88 KO, TRIF/MyD88 KO or TRIF/MyD88/STING KO cells showed no difference compared to WT control cells. KO of TRIF/MyD88 or TRIF/MyD88/STING in addition to MAVS did not increase infection rates further (Fig. 4C). To compare the IFN-stimulated response to YF17D and RIG-I ligand 3pRNA as a control stimulus in GM-DC/Mac between the genotypes CXCL10 mRNA expression was measured by qPCR. CXCL10 mRNA induction was reduced in MAVS KO and IFNAR KO GM-DC/Mac coinciding with the increase in viral replication, although the difference was not statistically significant (Fig. 4D). Collectively, these results indicate that YF17D is highly restricted by MAVS-dependent signaling downstream of RLRs in murine macrophages and DCs while MyD88 and TRIF-dependent signaling downstream of TLRs and the cGAS/STING pathway are not contributing to YF17D restriction.

**Figure 4.**
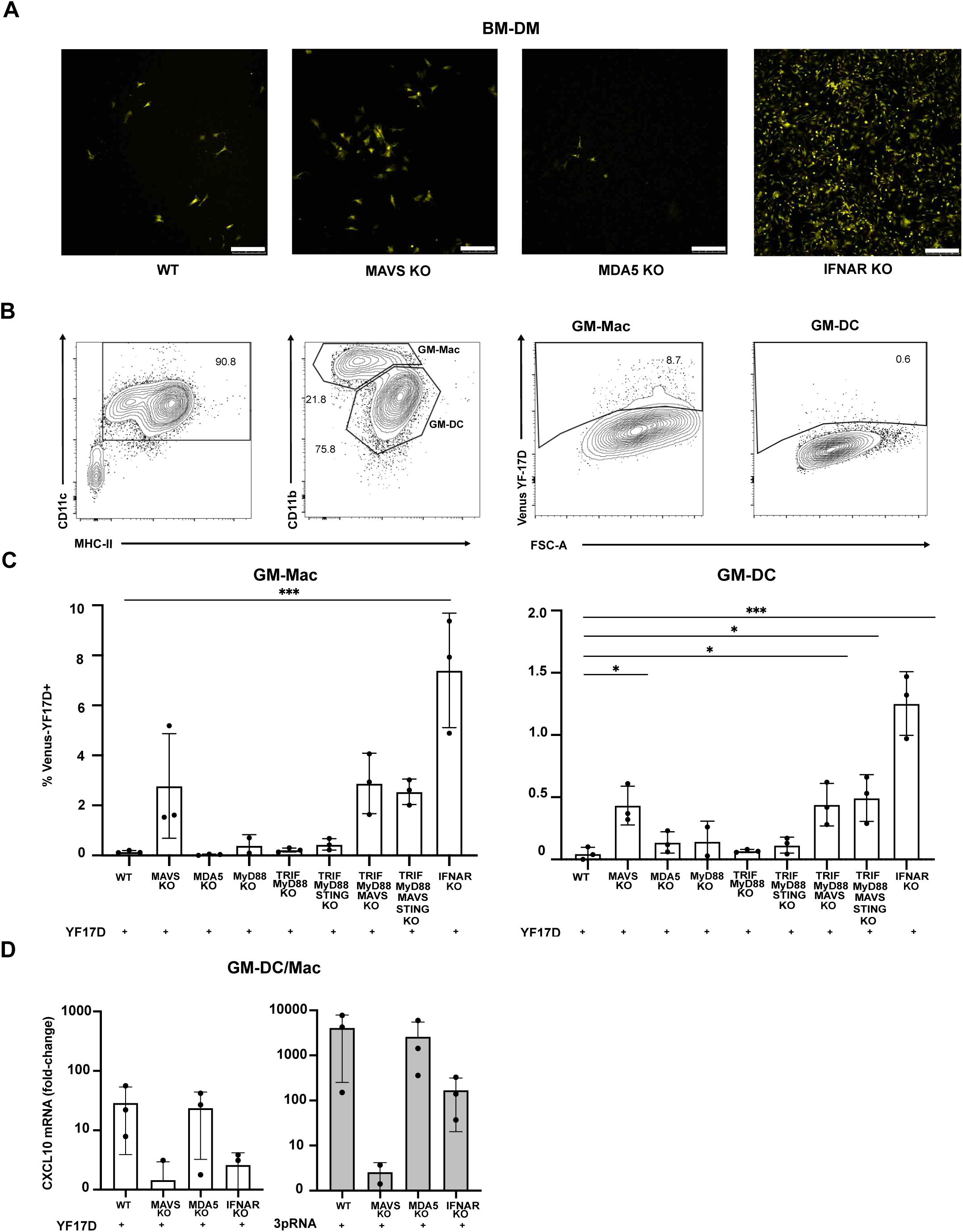
MAVS-dependent signaling restricts YF17D in murine macrophages and DCs. (A) Representative fluorescence microscopy images of BM-DMs infected with Venus YF-17D at 1 MOI for 48 hrs. Scale bars = 250 μm. (B-C) BM cells from the indicated knock-out mice were cultured with GM-CSF for 7 days and then infected with 5 MOI Venus-YF17D for 48 hrs. (B) Representative flow cytometry plots of the gating strategy for macrophages (GM-Mac) and DCs (GM-DC) from MAVS-KO. (C) GM-CSF-cultured BM cells from each indicated genotype were infected with 1 MOI Venus-YF17D. The percentage of Venus-positive cells in macrophages and DCs was measured by flow cytometry after 48 hrs. Mean and SD are shown in the bar graphs. One-Way ANOVA test with Dunnett’s correction was performed between WT and indicated knockouts (n=3, except for MyD88 where n=2). (D) GM-CSF-cultured BM cells from each indicated genotype were infected with 1 MOI Venus-YF17D or stimulated with RIG-I ligand 3pRNA. CXCL10 mRNA levels were quantified by qPCR after 24 hrs. Data were normalized to the relative expression of the HPRT reference gene and are shown as fold change relative to uninfected cells. One-Way ANOVA test with Dunnett’s correction was performed between WT and indicated knockouts. Error bars represent means and standard deviations (n=3).

To confirm these results in human cells we generated knockout cell clones by CRISPR/Cas9 gene editing using the 1205Lu human melanoma cell line which was previously shown to be responsive to RNA ligands of RIG-I, MDA5, and TLR3, but not TLR7/8 (30). After 48 hrs of infection the percentage of Venus^+^ infected cells was significantly increased in 1205Lu cells deficient in MAVS or RIG-I, but not in 1205Lu cells lacking MDA5 or TRIF (Fig. 5A and B). Double KO cell lines lacking MDA5/RIG-I, TRIF/MAVS, or TRIF/RIG-I showed an increased infection like that observed in single KO cell lines lacking MAVS or RIG-I or IFNAR. Thus, YF17D replication in 1205Lu cells was restricted by type I IFN and MAVS signalling with a major contribution of RIG-I. The increase of viral replication in MAVS and RIG-I KO cell lines was associated with a trend toward reduced YF17D-induced IFN-β mRNA expression. The strongest downregulation of IFN-β in response to YF17D was observed in MAVS and in RIG-I/MDA5 double KO cells. This reduction was comparable to that observed after stimulation with the RIG-I ligand 5’-triphosphate RNA that was used as a positive control. IFN-β mRNA expression after YF17D infection was also reduced in RIG-I single KO, but not to the same extent as in MAVS KO or MDA5/RIG-I double KO cells (Fig. 5C). IFN-β levels in the supernatants were below detection in YF17D-infected MAVS KO and MDA5/RIG-I double KO cells, but residual production of IFN-β was detected in RIG-I KO and MDA5 KO cells suggesting that MDA5 also contributes to the cumulative IFN-β response in 1205Lu cells (Fig. 5D).

**Figure 5.**
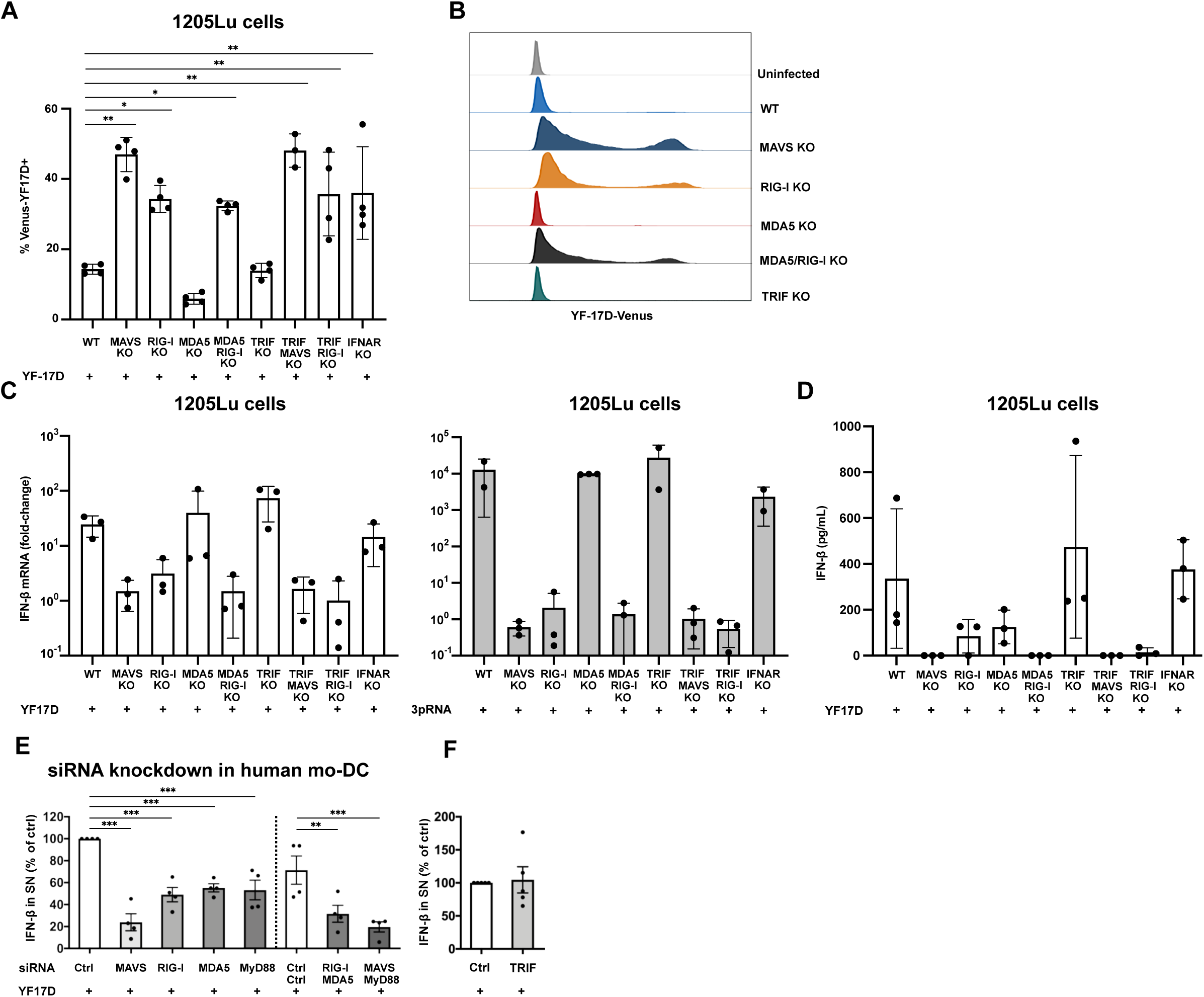
Sensing of YF17D in human cells requires RIG-I and MAVS. Wildtype (WT) and 1205Lu cells gene-edited by CRISPR-Cas9 mediated knockout (KO) for the indicated receptors were infected with Venus-YF17D at 5 MOI. (A) The percentages of Venus-YF17D positive cells were measured by flow cytometry after 48 hrs (mean +/- SD, n=4). Asterisks indicate statistically significant differences compared to WT control (**p<0.01, ***p<0.001, One-Way ANOVA test with Dunnett’s correction. (B) Representative histogram overlay plots showing the Venus fluorescence signal, from A. (C) 1205Lu cells of the indicated genotypes were infected with 5 MOI Venus-YF17D (left panel) or stimulated with RIG-I ligands 3pRNA (right panel). After 24 h, cells were collected for RNA extraction. IFN-β mRNA levels were quantified by qPCR. Data were normalized to the relative expression of the HPRT reference gene and are expressed as fold change relative to uninfected cells (mean+/- SD, n=3). (D) 1205Lu cells of the indicated genotypes were infected with 5 MOI Venus-YF17D and IFN-β concentrations in supernatants were measured by ELISA after 48 hrs (mean +/- SD, n=3). (C, D) One-Way ANOVA test followed by Dunnett’s correction was performed between WT and indicated knockouts; not significant. (E) Human Mo-DCs were transfected with siRNAs against the indicated target molecules or control siRNA. Double knockdown was performed with combined siRNAs (each 150 nM) and compared to 300 nM control siRNA. After 48 hrs mo-DCs were infected with 1 MOI YF17D for 36 hrs and IFN-β concentrations were measured in the supernatants by ELISA. Data points indicate results of independent experiments performed with mo-DCs from different donors. The data was normalized and is shown as percentage of the control siRNA condition (mean +/- SD, n=4). (F) In a separate set of experiments siRNA knockdown of TRIF was performed in mo-DCs as described in E. IFN-β concentrations in the supernatants were measured by ELISA 36 hrs post infection. Results are shown as percentages of the control siRNA condition (mean +/- SD, n=5). (E, F) Asterisks indicate statistically significant differences to the respective control siRNA condition (One-way ANOVA with Sidak correction **p<0.01, ***p<0.001).

Knockdown of MAVS using siRNA in primary human mo-DCs similarly showed that YF17D-induced IFN-β production is highly dependent on MAVS (Fig. 5E). Individual knockdown of RIG-I, MDA5, or MyD88 by siRNA also reduced IFN-β production, but not as strongly as MAVS knockdown (Fig. 5E, knockdown efficiency shown in Fig. S4). The combined knockdown of MAVS and MyD88 did not further reduce the IFN-β response. Knockdown of TRIF did not affect the IFN-β response to YF17D in mo-DCs (Fig. 5F), while the IFN-β response to LPS (positive control) was clearly reduced (Fig. S4). Thus, the IFN-β response of human DCs to YF17D infection also depends mainly on RLRs and MAVS-dependent signaling. Taken together, our results show that RLR/MAVS is the primary pathway sensing live YF17D in APCs and non-immune cells and that RIG-I is the dominant sensor.

### RIG-I interacts with 5’ phosphate dsRNA generated during YF17D infection

We sought to determine which viral RNA species are recognized by RIG-I. To this end, the 1205Lu parental cell line was infected with 1 MOI YF17D and the dsRNA structures formed were visualized by staining with a dsRNA specific antibody using confocal microscopy 24, 48 and 72 hrs after infection. dsRNA formation was observed in the cells’ cytoplasm after 48 and further increased after 72 hrs when the highest number of dsRNA positive cells was detected suggesting the formation of viral dsRNA intermediates (Fig. 6A). To further investigate the structural features required for the RLR-mediated response to YF17D infection, we isolated total RNA from wildtype 1205Lu cells 48 hrs after YF17D infection and digested it with RNA-modifying enzymes prior to transfection into WT 1205Lu cells (Fig. 6B). We observed that the immunostimulatory activity of RNA from infected cells (measured by CXCL10 induction) was lost after the treatment with RiboShredder and RNase III enzymes, which degrade total RNA and dsRNA, respectively. The RNase R enzyme, which specifically cleaves single-stranded RNA had no significant effect on CXCL10 induction compared to the control sample (no enzyme treatment). To test the requirement of 5’ phosphate residues for the activation of RLRs we treated the RNA with 5’-polyphosphatase (5’ PP) enzyme removing tri- and diphosphate groups from the 5’ end. CXCL10 induction by phosphatase-treated RNA was significantly decreased compared to induction observed with untreated RNA from infected cells (Fig. 6B). These results show that immunostimulatory RNA extracted from YF infected cells is double-stranded and contains a 5’ tri- or diphosphate groups, the classical characteristics of a RIG-I ligand.

**Figure 6.**
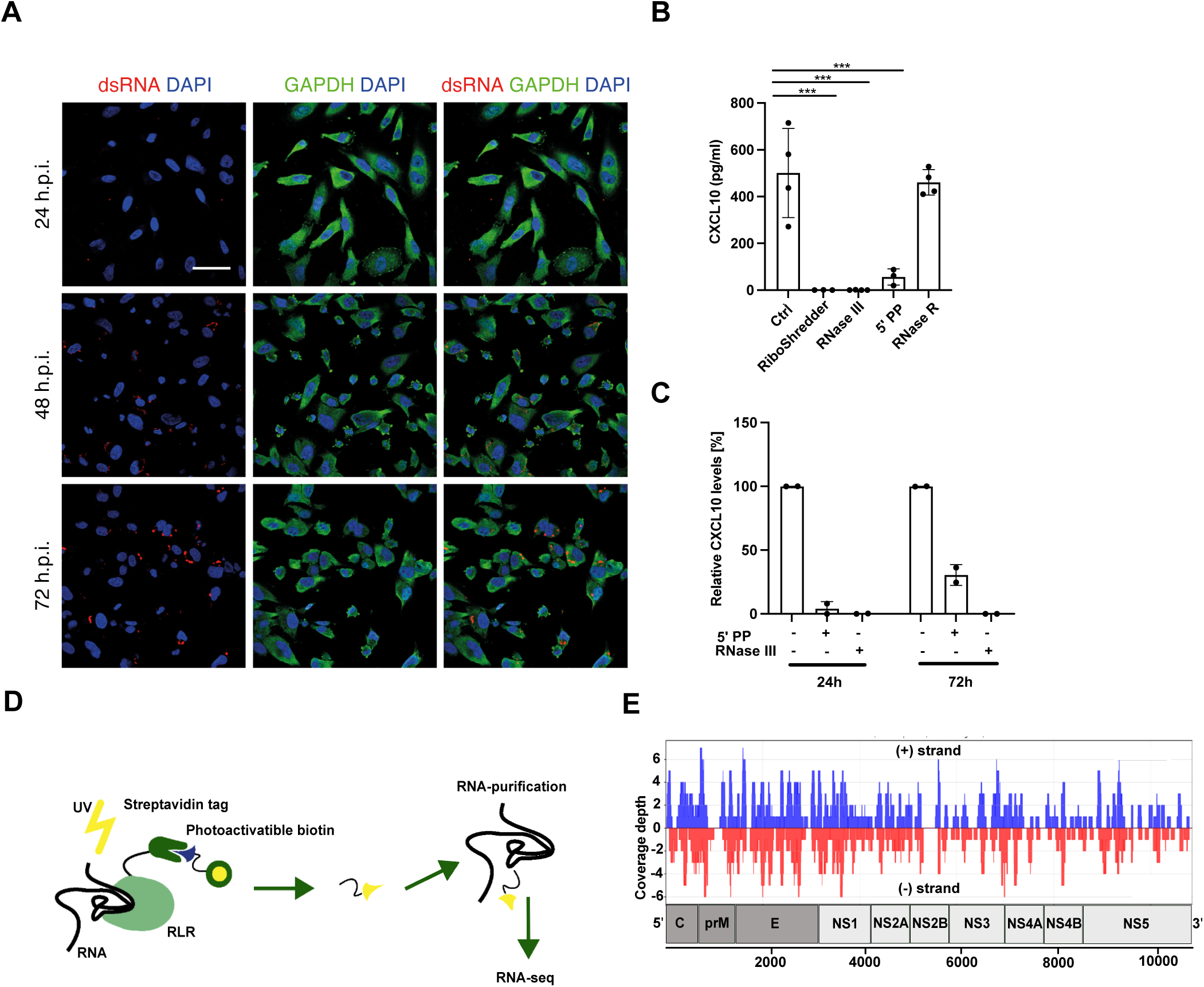
RIG-I interacts with 5’ phosphate dsRNA generated during YF17D infection. (A) Wildtype 1205Lu cells were infected with YF17D (MOI 1). At 24, 48,72 hrs p. i. cells were fixed, stained with anti-GAPDH (green), anti-dsRNA antibodies (red) and DAPI nuclear staining (blue) and analyzed by confocal microscopy. Representative images are shown. Scale bar, 50 µm, n=2. (B) RNA isolated from YF17D-infected 1205Lu cells at 48 hrs post infection was digested with the indicated enzymes and transfected into 1205Lu cells. CXCL10 concentrations were measured in the supernatants after 24 hrs by ELISA. One-Way ANOVA test with Dunnett’s correction was performed between control and treated samples (n=4, except RiboShredder condition where n=3; ***p<0.001). (C) RNA was isolated from YF17D-infected 1205Lu cells at 24 and 72 hrs post infection and treated with 5PP or RNAse III and then transfected into 1205Lu cells. CXCL10 levels were measured in the supernatants after 24 hrs. Results are shown as percentages of the untreated controls (w/o enzyme) from each experiment, n=2. (D) Schematic depiction of the lPL approach. (E) 1205Lu cells expressing N-mSAv-RIG-I were infected with 10 MOI YF17D for 48 hrs (one experiment). After infection, RIG-I/RNA complexes were isolated using streptavidin pull-down. The purified RNA was used to generate cDNA libraries, which were analyzed by strand specific NGS analysis. The read sequences were aligned to the YF genome reference in sense (blue) or antisense orientation (red). The y-axis depicts genome coverage; the x-axis represents positions on the reference genome.

We further evaluated whether the contribution of the 5’-phosphate and the double-strand structure to the immunostimulatory effect of RNA isolated from YF17D-infected 1205Lu cells differs between the early (24 hrs) and the late (72 hrs) time points. CXCL10 induction was significantly reduced after the digestion of dsRNA with RNase III at both time points post-infection compared to the untreated RNA control from the same time points (Fig. 6C). Enzymatic treatment with 5’-polyphosphatase (5’ PP) also reduced CXCL10 induction by RNA from infected cells at both time points. However, this effect was diminished at the later timepoint (Fig. 6C). Thus, induction of CXCL10 by RNA formed within 24 hrs of YF17D infection requires dsRNA with a 5’-phosphate end, pointing to a dominance of RIG-I ligand formation in the early phase after infection. At later time points, other PRRs such as MDA5 likely become more important

To test the hypothesis that RIG-I binds YF17D dsRNA intermediates we performed in *vivo* proximity labelling (IPL) followed by next-generation sequencing in 1205Lu cells at 48 hrs post infection with YF17D. As shown in the illustration (Fig. 6D) the IPL method relies on fusing a monomeric streptavidin (mSA) to a protein of interest (here RIG-I) and using a probe comprising a light-sensitive moiety and biotin to biotinylate proximal RNA upon UV irradiation. Biotinylation then captures the specific RNA and stabilizes the RNA-protein interaction ensuring that interacting RNAs are recovered after a streptavidin-based pull-down (31). To identify RIG-I–bound RNA of different polarities, we generated a strand-specific cDNA library. Biotinylated RNA isolated via streptavidin pulldown from infected 1205Lu cells was depleted of ribosomal RNA and sequenced. After removal of human-derived reads, which accounted for 82.7% of the total, 0.64% of reads in the RIG-I–tagged samples aligned to the YF17D reference genome, with an almost equal distribution between the forward (0.32%) and reverse (0.32%) strands (Fig. 6E). In the negative control sample with RNA purified from the streptavidin-based pulldown performed with infected 1205Lu expressing untagged RIG-I, only 0.064% of the reads were aligned to the YF17D genome. The equal distribution of aligned reads in positive and negative orientation indicates that YF17D dsRNA intermediates formed in infected cells interact with RIG-I.

## Discussion

In this study we showed that YF17D infects primary DCs and monocytes, whereas virus propagation is highly restricted by type I IFN receptor activation in these APCs. This limits the availability of endogenous viral antigen for presentation to T cells. We found that type I IFN induction and viral restriction are largely mediated by RLR/MAVS signaling and RIG-I is the dominant sensor interacting with YF17D dsRNA intermediates during YF17D replication.

Our observation that IFNAR blocking in human DCs and monocytes and IFNAR deficiency in murine DCs and macrophages significantly enhanced YF17D infection indicates that the autocrine or paracrine effect of type I IFN via IFNAR signaling is crucial for restricting YF17D replication and spread in APCs. Previous studies have shown that IFNAR-incompetent cell lines, such as BHK21 and Vero cells as well as IFNAR KO mice, are much more susceptible to viral infection with YF17D (22, 32, 33). Furthermore, individuals with IFNAR deficiencies were reported to suffer from severe adverse events after YF17D vaccination demonstrating the critical role of IFNAR signaling in controlling YF17D replication and limiting pathogenicity in vivo (7). The high sensitivity of YF17D to type I IFN seems to be related to the acquired mutations in the non-structural protein NS2A (34). Interestingly, using APCs generated from GM-CSF treated murine bone marrow (BM) cells we found that DCs were more resistant to YF17D infection than macrophages in the same culture even in the absence of IFNAR pointing to additional cell-intrinsic mechanisms for YF17D restriction in DCs. Limitations in viral entry and fusion may contribute to constitutive resistance of DCs as described previously for HIV and influenza virus in the cDC1 subpopulation (35).

We reported previously that the transcriptomic response of circulating DCs and monocytes is dominated by a global upregulation of IFN-stimulated genes and that peak expression of the costimulatory molecule CD86 correlates with plasma levels of IFN-induced CXCL10 (16) suggesting that type I IFN drives this activation. Consistently, IFNAR blocking experiments showed that autocrine and paracrine type I IFN play an important role for the activation of DCs and monocytes after infection with YF17D, but residual activation despite effective IFNAR-blockade indicated that direct effects of the virus or other cytokines also contribute. Our results are consistent with the critical role of type I IFN for differentiation, activation and maturation of DCs during viral infection and for the application of vaccines adjuvanted with immune stimulatory RNA like poly(I:C) (9, 11, 12, 14, 36, 37).

Despite reducing the activation of APCs, IFNAR-blockade increased the percentage of CD8^+^ T cells responding to restimulation by infectious YF17D in vitro without affecting the CD4+ T cell response. This differential effect was also observed in a vaccination cohort where peak IFN-α plasma levels correlated negatively with YF17D-responsive CD8^+^ T cells, but not CD4^+^ T cells. As type I IFN strongly restricts YF17D it also decreases the availability of viral antigens for presentation to T cells by APCs thereby limiting T cell activation. CD4^+^ T cells are more reliant on costimulation for their activation than CD8^+^ T cells (38). Therefore, the increased costimulation associated with type I IFN mediated activation of APCs likely counterbalances the effect of the reduced antigen dose on CD4^+^ T cell induction. The antigen dose seems to be critical for the CD8^+^ T cell response to YF17D vaccination as the extent of YF17D viremia determines the magnitude of CD8^+^ T cell induction (2). Our results suggest that after yellow fever vaccination the limiting effect of type I IFN on antigen dose is prevailing over direct effects of type I IFN promoting CD8^+^ T cell survival, expansion and differentiation (39).

In line with our finding, transient local blockade of type I IFN signaling can enhance the immunogenicity of several viruses and live viral vaccines (including YF17D) in mice leading to persistently higher frequencies of specific T cells and higher antibody titers (17, 40). Blocking IFNAR temporarily during infection and vaccination may have additional benefits. For example, it was shown recently in the mouse model that short term IFNAR blockade during acute viral infection and mRNA-LNP vaccination promotes differentiation of stem cell-like memory CD8^+^ T cells mediating long-term protection (41).

A well-controlled type I IFN response is critical for balancing safety and efficacy of the yellow fever vaccine and likely also other live viral vaccines, preventing pathology due to virus dissemination while allowing sufficient production of viral antigens. Given the importance of controlling YF17D replication, we investigated the mechanism of YF17D sensing and restriction. RLR/MAVS signaling was found to be crucial for inducing the type I IFN response and restricting YF17D in immunocompetent cells including primary DCs and macrophages with a predominance of RIG-I over MDA5. This is consistent with previous reports of RIG-I/MAVS-dependent type I IFN induction in non-immune cell lines and in infected pDCs (23, 25, 27). By combined KO of signaling adaptors, we could show that neither MyD88- nor TRIF-dependent TLRs nor cGAS/STING contributed to YF17D restriction in murine DCs and macrophages. According to our results, MyD88-dependent TLR signaling may have a minor contribution in human mo-DCs, but the RIG-I/MAVS pathway was clearly more important for type I IFN induction in these cells. In contrast, Querec et al. described TLR/MyD88-dependent activation of murine splenic DCs (22), which may be explained by the absence of viral replication and additionally by the presence of pDCs within the murine splenic DC fraction as pDCs have been shown to sense immature YF17D particles and viral RNA transferred from adjacent infected cells via TLR7 (23, 24). Reactive oxygen species (ROS) were shown to contribute to the type I IFN response in human mo-DCs demonstrating the importance of cellular metabolism and mitochondrial function for RLR/MAVS-mediated YF17D sensing (27). We found that YF17D infection leads to the formation of dsRNA intermediates with 5’-phosphate groups as ligands for RIG-I. It has previously been shown that RIG-I exhibits a preference for short dsRNA with a 5’-di or triphosphate group, whereas MDA5 predominantly binds to long double-stranded RNA (9, 42–44). We observed that the requirement for a tri- or diphosphate moiety at the 5’ end became less stringent at later time points after infection suggesting that dsRNA without a 5’- tri- or diephosphate end is formed later and may contribute to the response, e.g., by triggering MDA5 activation. Such sequential sensing by RIG-I and MDA5 has been reported for West Nile Virus infection (45). It was also shown that the type I IFN response to Dengue virus is mediated by sensing of full length 5’-tri-phoshpate-dsRNA by RIG-I with additional contribution of MDA5 (46). Thus, RLR/MAVS activation by replicative dsRNA seems to be a common sensing mechanism for several orthoflaviviruses.

Taken together our study shows that the safety and efficacy of the live attenuated vaccine YF17D relies on a closely regulated type I IFN response triggered by RLR-dependent sensing of replicative viral dsRNA.

## Methods

### Virus stocks

For functional assays, two yellow fever virus variants were produced and purified: YF17D, derived from a Stamaril vaccine dose (Sanofi), and Venus-YF17D, a fluorescent reporter virus. The Venus-YF17D plasmid was kindly provided by Charles M. Rice and Margaret MacDonald (The Rockefeller University, New York, USA). Virus production was performed in BHK-21 cells (DSZM, ACC61) or Vero B4 cells (DSMZ, ACC33) and followed standard protocols (47, 48). BHK-21 or Vero B4 cells were cultured in DMEM (Sigma Aldrich) with 10% FBS (Gibco), 2 mM L-glutamine, 1% Pen/Strep (Thermo Fisher Scientific) and infected at an MOI of 0.1. Supernatants were collected upon visible cytopathic effect, clarified, and mixed with 7% (w/v) PEG 8000. After incubation at 4°C, viral particles were pelleted, resuspended in TNE buffer (20 mM Tris-HCl pH 8, 150 mM NaCl, 2 mM EDTA), and purified via sucrose gradient (30% and 60%) ultracentrifugation. Final virus stocks were stored in TNE buffer and quantified by plaque assay. Plaque assays were performed on BHK-21 or Vero B4 cells to determine YF17D and Venus-YF17D titres. Serial ten-fold virus dilutions in Opti-Mem were added to duplicate wells. After 1 hr incubation at 37 °C, the inoculum was removed and replaced with an agarose overlay (DMEM, 10% FBS, L-glutamine, Pen/Strep). After 3 to 5 days of incubation the agarose overlay was removed, and cells were stained with 2.5% crystal violet. Plaques were counted, and titres expressed as PFU/mL.

### YF17D vaccination cohort

A total of 250 flavivirus-naïve young adults were recruited at LMU Munich (2015–2019). All participants received 0.5 mL YF17D (Stamaril) subcutaneously, and peripheral blood samples were collected at baseline and on days 3, 7, 14, and 28 post vaccination. The study was approved by the LMU ethics committee (IRB #86-16) and registered at ISRCTN (ISRCTN17974967) (49).

### Human PBMC and monocyte isolation

PBMC from healthy donors were isolated either from leucocyte reduction chambers obtained from thrombocyte donations (LMU Klinikum, Munich, Germany) or from freshly drawn peripheral blood of healthy donors who had given informed consent (with approval of the LMU ethics committee, no. 18-415). Blood was drawn using S-Monovette Sodium-Heparin (Sarstedt). PBMC were isolated by gradient centrifugation using Biocoll (Biochrom), washed once with PBS, and red blood cells were lysed by incubating with red blood lysis buffer (Sigma Aldrich) for 10 min at RT. The PBMCs were subsequently washed twice using PBS with 2 mM EDTA and 2% FCS. Total monocytes were isolated by magnetic-activated cell sorting (MACS) using a Classical Monocyte Isolation Kit or Pan Monocyte Isolation Kit (Miltenyi Biotec). For generation of mo-DCs, monocytes were positively selected by MACS using human CD14 microbeads (Miltenyi Biotec) according to manufacturer’s protocol (purity > 95%).

### Generation of human mo-DCs from CD14^+^ monocytes

CD14^+^ Monocytes were cultured at 2×10^6^ cells per well in a 6-well plate in 2 ml DC medium (RPMI 1640, Biochrom, 10% FCS, Biochrom, 100 U/mL Pencillin, 100 µg/mL Streptomycin, 1% NEAA, 2mM GlutaMAX, 1mM Na-pyruvate (Thermo Fisher Scientific), 0.05 mM β-mercaptoethanol, Sigma Aldrich) containing 500 U/mL GMCSF (Peprotec) and 500 U/mL IL-4 (Invivogen). In smaller well-sizes, the same concentration of 10^6^ cells/mL medium was used. Cells were incubated at 37 °C for 4 days, and on day 4, IL-4 was resupplied at 250 U/mL.

### Culture and infection of human PBMC, mo-DCs and monocytes

PBMC, mo-DCs and monocytes were infected with different MOIs of purified Stamaril-derived YF17D or Venus-YF17D by adding the respective amount of virus to the medium and culturing for 36 (mo-DCs) or 48 hrs (PBMC and monocytes). During the infection mo-DCs were cultured in DC medium containing GM-CSF (500 U/mL) and IL-4 (500 U/mL) and monocytes were cultured in DC medium containing low-dose GMCSF (200 U/mL) for 48 hrs. To inhibit the type I IFN response, polyclonal rabbit anti-IFN-α antibody (PBL31101-1), rabbit anti-IFN-β (PBL21385-1) from PBL Assay Science, and mouse monoclonal anti-IFNAR2 antibody (PBL31410-1, PBL Assay Science, clone MMHAR-2, Acris) were used at concentrations of 2 000 U/mL, 5 000 U/mL, and 10 μg/mL respectively. The antibody cocktail was added to the cells 30 min prior to infection. Lipopolysaccharide (LPS) from *E. coli* (L4391, Sigma-Aldrich) was used at a concentration of 10 ng/mL, R848 (Invivogen) was used at 3 µM.

### SiRNA knockdown in human mo-DC

Monocytes were cultured in 24-well culture plates at 5% CO2, 37℃ in DC medium at a density of 1×10^6^ or 0.5×10^6^ cells/mL in a volume of 0.5 mL depending on the experiment. 500 U/mL of recombinant human GM-CSF (Peprotech, USA) and 500 U/mL IL-4 (Invivogen, USA) were added and mo-DCs were harvested on day 4. DCs were electroporated with ON-TARGETplus SMARTpool siRNAs against RIG-I (M-012511-01), MDA5 (L-013041-00-0005), MAVS (M-024237-02), MyD88 (L-004769-00-0005) or TRIF/TICAM (L-012833-00-0005) or with control non-targeting siRNA (D-001810-10-05) from Dharmacon/Horizon Discovery using the Neon Transfection System (Thermo Fisher). Briefly cells were washed in PBS twice and resuspended in Buffer T at 2 x 10^7^/mL mixed with individual siRNA pools at previously optimized concentrations (150-200 nM) or two siRNA pools (each 150 nM) and electroporated (10 µl tip, 1500V, single pulse, 20 ms). Cells were immediately added to 500µl prewarmed DC medium without antibiotics (2 x 10^5^/24-well). Fresh GM-CSF and IL-4 (500 U/mL each) were added and mo-DCs were infected with 1 MOI YF17D for 36 hrs. The IFN-β concentration in the supernatants was measured by ELISA (IFN-β DuoSet, R&D Systems, USA) following the manufacturer’s protocol.

### Culture and infection of 1205Lu cells

Wildtype 1205Lu cells were provided by Dr. R. Besch (LMU Munich). The 1205Lu knockout cells were generated by CRISPR-Cas9 gene editing as described (50). Cells were cultured in DMEM (Sigma-Aldrich) containing 10% FCS, 1% L-glutamine, and 100 U/mL penicillin and streptomycin under standard conditions (37°C, 5% CO2, >90% humidity). CRISPR/Cas9 knockout or wildtype human melanoma 1205Lu cells (2.5 × 10□ cells/well) were infected with Venus-YF17D virus, which was diluted to the desired multiplicity of infection (MOI) in Opti-MEM (Gibco, Thermo Fisher Scientific). Transfection of the RIG-I ligand 3pRNA (0.5 mg/mL) into 1205Lu cells was performed using 0.3 μL Lipofectamine RNAiMAX (Thermo Fisher Scientific) following the manufacturer’s instructions. The 5’-triphosphate RNA was generated in vitro as described previously (50). Cells were incubated for 24 hrs for isolating RNA and performing qPCR or for 48 hrs for detecting infected cells by flow cytometry and for collecting the supernatants to measure CXCL10 by ELISA.

### Isolation, differentiation, and infection of murine BM-derived DCs

Wildtype and knockout (KO) C57BL/6 mice, including knockout for MyD88, TRIF/MyD88, TRIF/MyD88/MAVS, TRIF/MyD88/STING, and TRIF/MyD88/STING/MAVS, were kindly provided by Prof. Ulrich Kalinke (TWINCORE, Hannover, Germany). All mice were bred and genotyped under pathogen-free conditions in the animal facility at the University Hospital, LMU. Male or female mice aged between 6 and 16 weeks were used. BM cells obtained from the femurs and tibiae of C57BL/6 mice were cultured in 10 cm-dishes. For generation of GM-CSF-derived DCs and macrophages (GM-DC/Mac) 2×10^5^ cells/mL BM cells were seeded in 10mL RPMI 1640 medium (Sigma-Aldrich) with 10% FCS, 1% L-glutamine, 1% Pen-Strep, 1% NEAs, 1% Na-Pyruvate and 20 ng/mL murine GM-CSF (Peprotech). For microscopy analysis, BM-DM were generated by culturing BM cells at 5 ×10^5^ cells/mL in 10ml DMEM with 10% FCS, 5% horse serum, 1% Pen-Strep, 1% Na-Pyruvate and 20 ng/mL murine M-CSF. DCs and macrophages were cultured at 37°C in a humidified atmosphere with 5% CO_2_ for 7 days. The differentiating medium containing growth factors was freshly added to 20mL and then exchanged every 2 days for macrophages or 3 days for DCs. The DC and macrophage phenotypes were confirmed by flow cytometry. GM-DC/Mac were harvested and seeded in a 12-well plate (5×10^5^ cells/well) for FACS analysis and in a 96 well plate (5×10^4^ cells/well) for qPCR. M-CSF-derived macrophages were seeded in a 96 well plate (2×10^4^ cell/well) for fluorescence microscopy. Cells were infected with Venus-YF17D (MOI 1 or 5). After 24 hrs, cells were lysed and CXCL10 levels were detected by qPCR as described below. After 48 hrs, YF17D infection was detected by monitoring the Venus expression level by fluorescence microscopy and by flow cytometry.

### T cell stimulation with YF17D virus

To measure antigen-specific T cells, cryopreserved PBMC were thawed and rested overnight at high density (5×10^6^ million cells per mL) in R10 medium at 37°C in a 5% CO2 humidified atmosphere. Cells were infected with 1 MOI of Venus-YF17D virus or with the equivalent volume of purified supernatant of uninfected cells (unstimulated control). In the condition with IFNAR blockade, α-IFNAR (clone MMHAR-2) was added 30min before the virus at 2.5µg/mL. After 24 hrs Brefeldin A (BioLegend) was added for an additional 20 hrs.

### Flow cytometry

Antibodies used for flow cytometry are listed in Supplementary Table S1. Human PBMC, mo-DCs and monocytes (Fig. 1 and 2) were harvested, transferred onto a 96 well plate, washed and resuspended in PBS with 2% FCS and 2mM EDTA. Cells were stained with fluorescently labelled antibodies (see Table S1) for 30 min at 4 °C washed twice and then fixed with BD Cytofix (BD Biosciences). For intracellular E protein staining cells were permeabilized using the eBioscience Foxp3/Transcription Factor staining kit (ThermoFisher Scientific) and stained with AlexaFluor 647-conjugated 4G2 antibody (D1-4G2-4-15, Novus Biologicals) at a dilution of 1:100. After the T cell re-stimulation with YF17D (Fig. 3), PBMCs were stained with fixable Viability Dye eFluor™ 780 (Thermo Fisher Scientific) and then blocked for 10 min with 10 % human AB serum (Sigma Aldrich). Surface staining was performed in blocking buffer for 20 min. For intracellular cytokine staining (ICS) cells were fixed and permeabilized with Foxp3/Transcription Factor Staining Buffer Set (Invitrogen) for 20 min at RT and incubated with antibodies against IFN-γ, TNF-α, CD40L in permeabilization buffer for 45 min at RT. The percentages of CD40L^+^IFNγ^+^ CD4^+^ T cells and IFNγ^+^TNFα^+^ CD8^+^ T cells obtained in the unstimulated sample were subtracted from those in the YF17D-stimulated sample. Murine GM-Mac and GM-DCs cells (Fig. 4) were harvested, transferred onto a 96 well plate, washed in PBS with 0.5% BSA and 2mM EDTA and incubated in FcR blocking reagent TruStain fcX (BioLegend) for 10 min at RT. Cells were then stained with antibodies against cell surface molecules and with LIVE/DEAD fixable Near-IR stain (Thermo Fisher Scientific) for 15□min at RT. Cells were washed and fixed in 4% PFA at RT for 20 min. Samples were measured using the BD LSRFortessa (BD Biosciences) or the CytoFLEX S (Beckman Coulter) or Cytek Aurora (Cytek Biosciences). Data was analyzed using FlowJo software v9 and v10.

### ELISA

Cytokine levels in the cell culture supernatant were determined using the commercial IFN-β DuoSet ELISA kit (R&D Systems) or Human IP-10 (CXCL10) ELISA Set (BD Biosciences) according to the manufacturer’s protocols.

### IFN-α detection in human plasma

IFN-α concentrations in the plasma if YF17D vaccinees were measured by PBL Assay Services (PBL Assay Science) using the MSD S-Plex human IFN-α2a kit.

### Reverse transcription PCR (RT-PCR) and quantitative real-time PCR (qPCR) analysis

Total cellular RNA from cell lysates was extracted using the PeqGOLD kit (Peqlab). RNA was reverey transcribed into cDNA with random hexamers using the RevertAid First-Strand Synthesis System for RT-PCR (Thermo Fisher Scientific). The gene-specific human or murine primers were designed using the universal probe library canter (Roche). huIFNb: (forward) CGACACTGTTCGTGTTGTCA, (reverse) GAGGCACAACAGGAGAGCAA; huHRPT: (forward) TGACCTTGATTTATTTTGCATACC, (reverse) CGAGCAAGACGTTCAGTCCT; muCXCL10: (forward) GCTGCCGTCATTTTCTGC, (reverse)TCTCACTGGCCCGTCATC; muHRPT: (forward) GGAGCGGTAGCACCTCCT, (reverse) CTGGTTCATCATCGCTAATCAC. qPCR was performed using the Roche Light Cycler 480-II/96 system. Data were normalized relative to the expression of the HPRT reference gene. The 2^-△△CT method was used for gene expression analysis and to calculate the fold-changes by normalizing to uninfected control.

### Immunofluorescence and confocal microscopy

1205Lu wildtype cells were seeded onto coverslips in 24-well plates at the density of 5×10^4^ cells/well in 1mlL DMEM culture medium and incubated at 37° C for 24 hrs. Prior to seeding the coverslips were cleaned by incubation in 1 M hydrochloric acid (HCl) overnight, washed thoroughly with water, and stored in 70% ethanol until use. The cells were then infected with 1 MOI YF17D. After 24, 48, and 72 hrs the medium was removed, the cells were washed with PBS and fixed with 4% PFA for 30 minutes at 37°C followed by permeabilization with 0.3% Triton X-100 (in PBS) for 5 minutes, and blocking with PBS containing 0.3% Triton X-100 and 3% goat serum for 1 hr at RT. Cells were stained with primary antibodies against dsRNA (J2, Scicons) and GAPDH (FL-335, Santa Cruz Biotechnology, 1:1000) at 4°C for 24 hrs in blocking buffer and washed 3x with PBS containing 0.3% Triton X-100. Cells were incubated with secondary antibodies (goat anti-mouse IgG Alexa Fluor 647 and IgG Alexa Fluor 488 (Invitrogen, Thermo Fisher Scientific) diluted 1:500 in blocking buffer, for 1 hr at RT. After 2 washes with PBS/0.3% Triton X-100, cells were stained with 4’,6-diamidino-2-phenylindole (DAPI) 1:1000 in PBS for 3 min, washed 3 times with PBS and applied to slides with 20 μL of Mowiol 4-88, mounting medium (Carl Roth). Confocal microscopy was performed using the Leica TCS SP5 microscope.

### Enzymatic treatment and re-transfection of RNA

Wildtype 1205Lu cells were seeded in a 6-well plate at 2×10^5^ cells/well and infected with YF17D virus at 3 MOI in 2 mL DMEM culture medium. After 24, 48 or 72 hrs total RNAs were extracted using TRIzol reagent (Invitrogen, Thermo Fisher Scientific) following the manufacturer’s instructions. The extracted RNAs (1 µg/condition) were treated with RiboShredder (Epicentre, Madison), RNase III (Thermo Fisher Scientific), RNase R (Epicentre, Madison), 5’-polyphosphatase (Biosearch Technologies) or FastAP (Thermo Fisher Scientific) for 30 minutes at 37°C. 1205Lu cells seeded in a 96-well plate (2.5 ×10^3^cells/well) were transfected with 100ng of each RNA using 0.3 μL Lipofectamine RNAimax (Thermo Fisher Scientific) following the manufacturer’s instructions. The concentration of CXCL10 in the supernatants was measured by ELISA after 24 hrs.

### In *vivo* proximity labelling (IPL) and next generation sequencing (NGS)

1205Lu cells (17×10^6^) expressing human RIG-I fused to a monomeric streptavidin tag were seeded on a 500 cm^2^ plate and infected with 10 MOI YF17D for 48 hrs. Biotin-TFPA-PEG3 (Thermo Scientific) 1 μM was added and cells were incubated for 2 hrs at 37°C. Cells were washed twice with ice-cold PBS. For photolabeling, 8 mL of cold PBS was added to the cells, and they were irradiated under UVA light (312 nm) at 500 mJ/cm² for 1 minute. After cross-linking, biotinylated cells were spun down and lysed, and total RNA was extracted using the phenol-chloroform extraction method. The isolated RNA was incubated with streptavidin beads for 4 hrs at 4°C, and the specific RIG-I/RNA complexes recovered by streptavidin pull-down were checked for quality using an RNA 6000 Pico chip (Bioanalyzer 2100, Agilent) and analyzed by NGS. After RNA isolation using TRIzol reagent (Invitrogen), rRNA was removed from the samples (NEBNext rRNA Depletion Kit, New England Biolabs) and cDNA libraries were generated using the NEBNext Ultra II Directional RNA Library Prep Kit for Illumina (New England Biolabs) following the manufacturer’s instruction. The quality of cDNA libraries was evaluated using a High Sensitivity DNA Chip (Bioanalyzer 2100, Agilent). Sequencing was performed using the Illumina NextSeq 500 platform. Raw sequencing reads were trimmed to remove adapters and low-quality bases using Trim Galore v.0.6.5 with default parameters (51). Filtered paired-end reads were aligned in two steps using Bowtie2 v.2.5.0 with default settings (52). SAM output files from Bowtie2 were converted to BAM format, sorted, and indexed using SAMtools v.1.21 (53). In the first alignment step, reads were mapped to the human reference genome (hg38) was performed to identify non-viral reads. The remaining unmapped reads were subsequently aligned to the Yellow Fever virus 17D reference genome (GenBank accession NC_002031.1). Genome-wide coverage profiles for every nucleotide of the YF17D genome were generated using SAMtools depth function.

### Statistical analysis

All statistical analyses were performed using GraphPad Prism (version 9). For comparisons across multiple groups, either a non-parametric Kruskal–Wallis test followed by Dunn’s post-hoc correction or a one-way ANOVA with Dunnett’s correction was applied, as indicated in the figure legends. Paired Wilcoxon matched pairs signed rank tests were used for matched donor samples. Correlation analyses were performed using Spearman’s rank correlation.

## Acknowledgements

The authors thank Yvonne Schäfer and Sandra Reiner for technical assistance. The authors also thank Arne Kroidl, Günter Fröschl and Kristina Huber for serving as clinical study investigators. We acknowledge the Core Facility Flow Cytometry of the Biomedical Center, LMU Munich, and thank Lisa Richter, Pardis Khosravani and Benjamin Tast. The authors also thank all the cohort participants who voluntarily participated in the study and donated samples. Parts of this work have been performed for the doctoral theses of MZ, EW, AD, ASP, PS, GS, MKS, LR at the LMU Munich. The project was funded by a combined grant of the German Research Foundation (DFG) project no. 391217598 to SR and ABK and the French National Research Agency (ANR) project no. ANR-17-CE15-0031-01 to GBS and by the DFG project no. 369799452 TRR237-TPB14 to ABK. and SR The project has received additional funding from the European Union’s Horizon Europe research and innovation program under Grant Agreement no. 101137459 - Yellow4FLAVI – “Deconstructing the protective immunity of yellow fever virus 17D to inform flavivirus vaccine design” to SR, ABK and GBS. Grants of the iMed consortium of the German Helmholtz Societies and the Einheit für Klinische Pharmakologie (EKLIP), Helmholtz Zentrum München, Neuherberg, Germany to SR further supported this work. EW and JTS received funding from the Friedrich-Baur-Stiftung and EW received a scholarship from the Villigst Foundation. PS, GS and MKS received funding from the FöFoLe Program of the Medical Faculty of the LMU Munich, MZ, ASP, GS, and MKS from the international doctoral program “iTarget: Immunotargeting of cancer” funded by the Elite Network of Bavaria and MZ and ASP from the doctoral program iTarget 2.0.

## Author contributions

MZ, EW, AD, ASP designed and performed experiments, analyzed and interpreted data, and contributed to writing the manuscript. PS, JTS, FD, GS, MKS, LR, YH, VA designed and performed experiments and analyzed data. JTS, KE and HK contributed to experimental design and data interpretation. MP and JTS helped to initiate the cohort, MP performed vaccinations and served as clinical study investigator. JS and UK contributed mouse models. GBS contributed unique reagents and acquired funding. ABK and SR conceptualized the study, contributed to experimental design, interpreted data, acquired funding, supervised and wrote the manuscript. All authors critically reviewed and approved the manuscript.

