## Supplementary Information for "RIG-I-like receptor-dependent type I Interferon regulates antigen dose and activation in yellow fever vaccine 17D-infected antigen presenting cells"

List of Supplementary Figures

Figure S1

Figure S2

Figure S3

Figure S4

Supplementary Table S1

Supplementary Methods

### Supplementary Figures

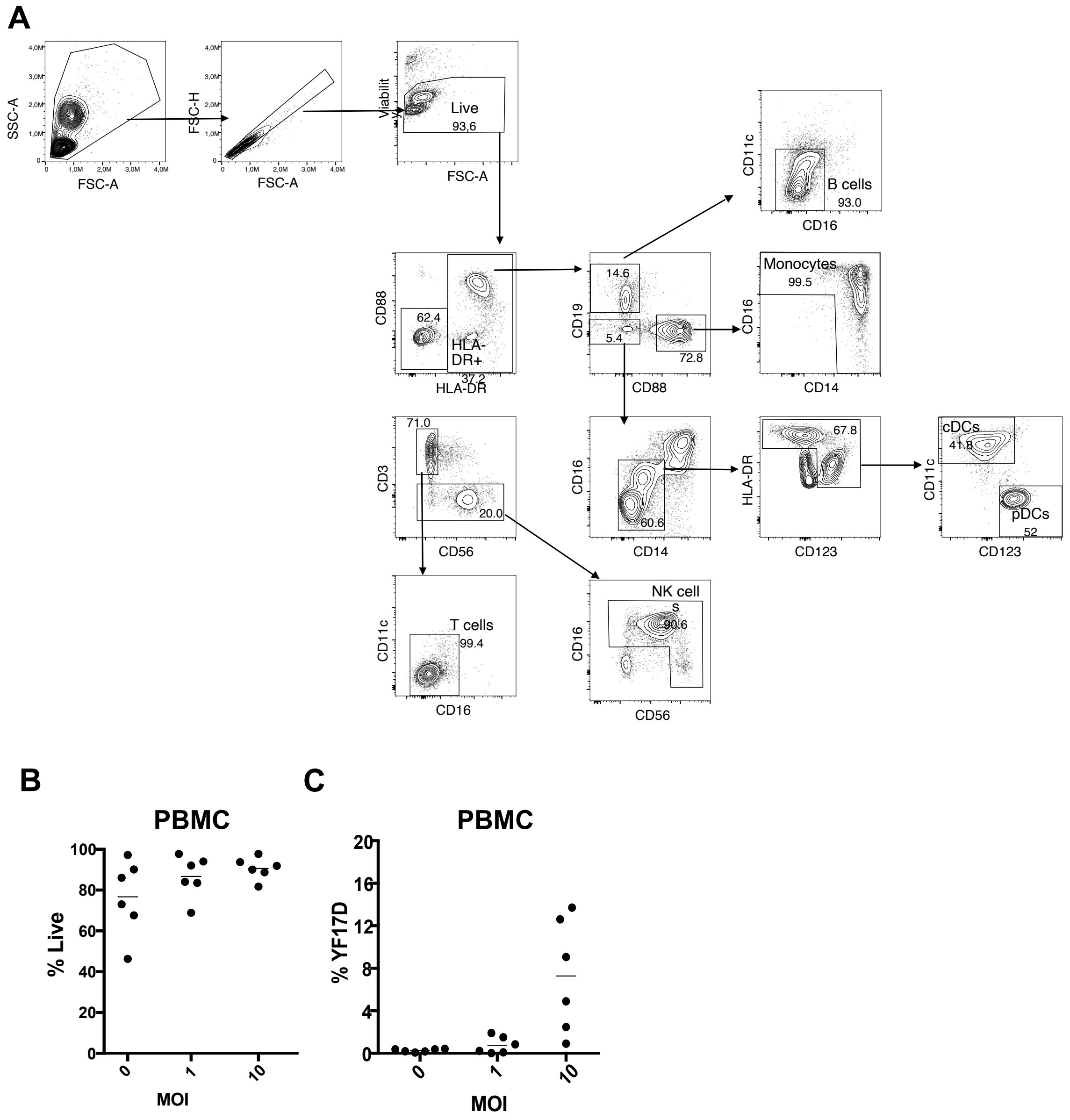


**Figure S1 Flow cytometric analysis of YF17D infection in PBMC**

(A) Gating strategy in PBMC (for data shown in Figure 1A. (B) Left panel: Percentage of living cells as determined by life/dead staining in PBMC after 48 hrs infection with 1 or 10 MOI compared to the uninfected condition. Right: Percentage of cells positive for E protein measured by flow cytometry after intracellular staining using 4G2 antibody.

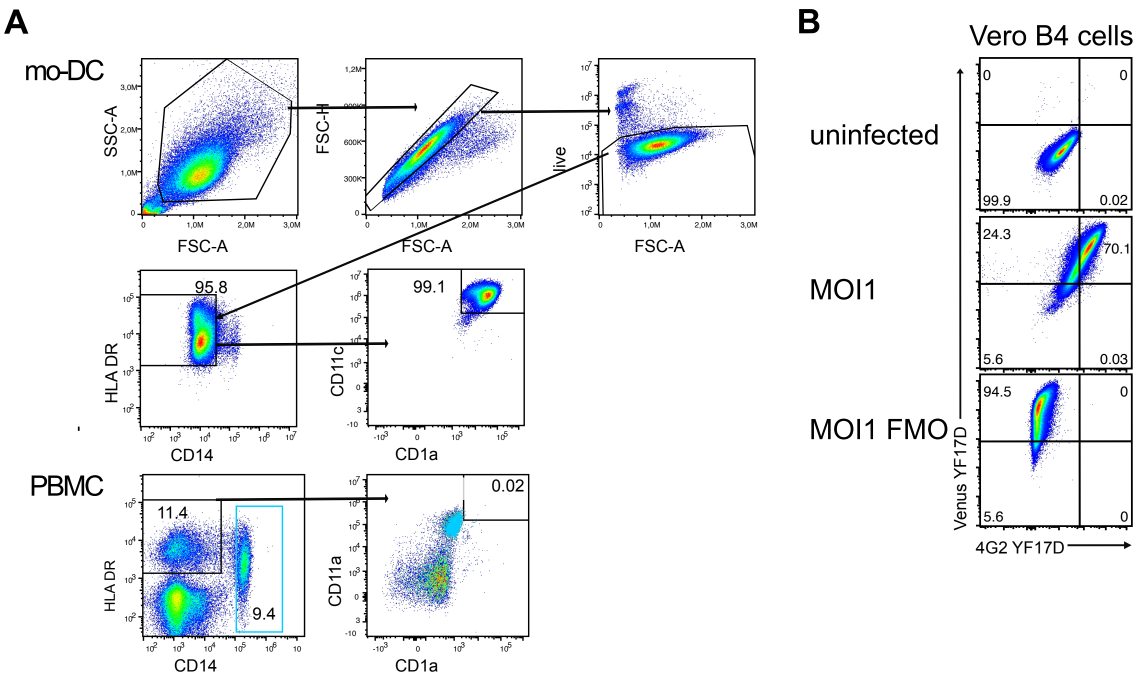


**Figure S2 Gating strategy for mo-DCs and detection of E protein in Venus-YF17D infected cells**

(A) Gating strategy for mo-DCs. PBMC stained for the same markers are shonw for comparison to demonstrate the change in phenotype after mo-DC generation from monocytes. (B) Vero B4 cells were infected with Venus-YF17D at 1 MOI or not infected and subsequently stained intracellularly with 4G2 antibody to detect the viral E protein. Representative results are shown as pseudocolor dot plots. Quadrant gates were set according to the uninfected control. Infected cells not stained with 4G2 antibody are shown in the bottom plot as fluorescence minus one (FMO) control.

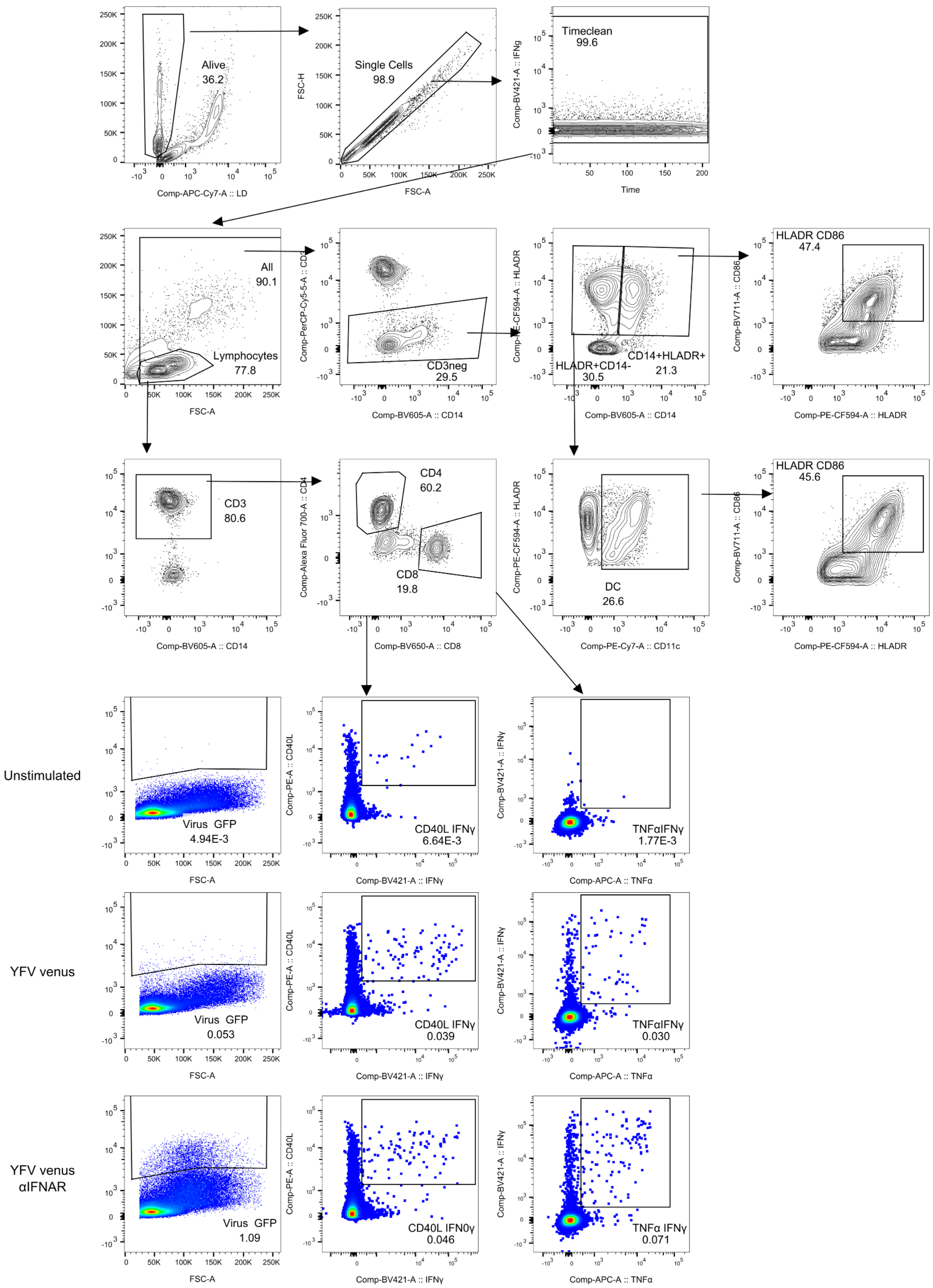


**Figure S3 Gating strategy for PBMC data shown in Figure 6**

The gating strategy for monocytes, sDCs and T cells in PBMC after restimulation with Venus-YF17D is shown in the upper panels. Below, representative results of the Venus fluorescence (left), CD40L and IFN-γ detection in CD4+ T cells (middle) and TNF-α and IFN-γ detection in CD8+ T cells (right) for the indicated conditions are shown.

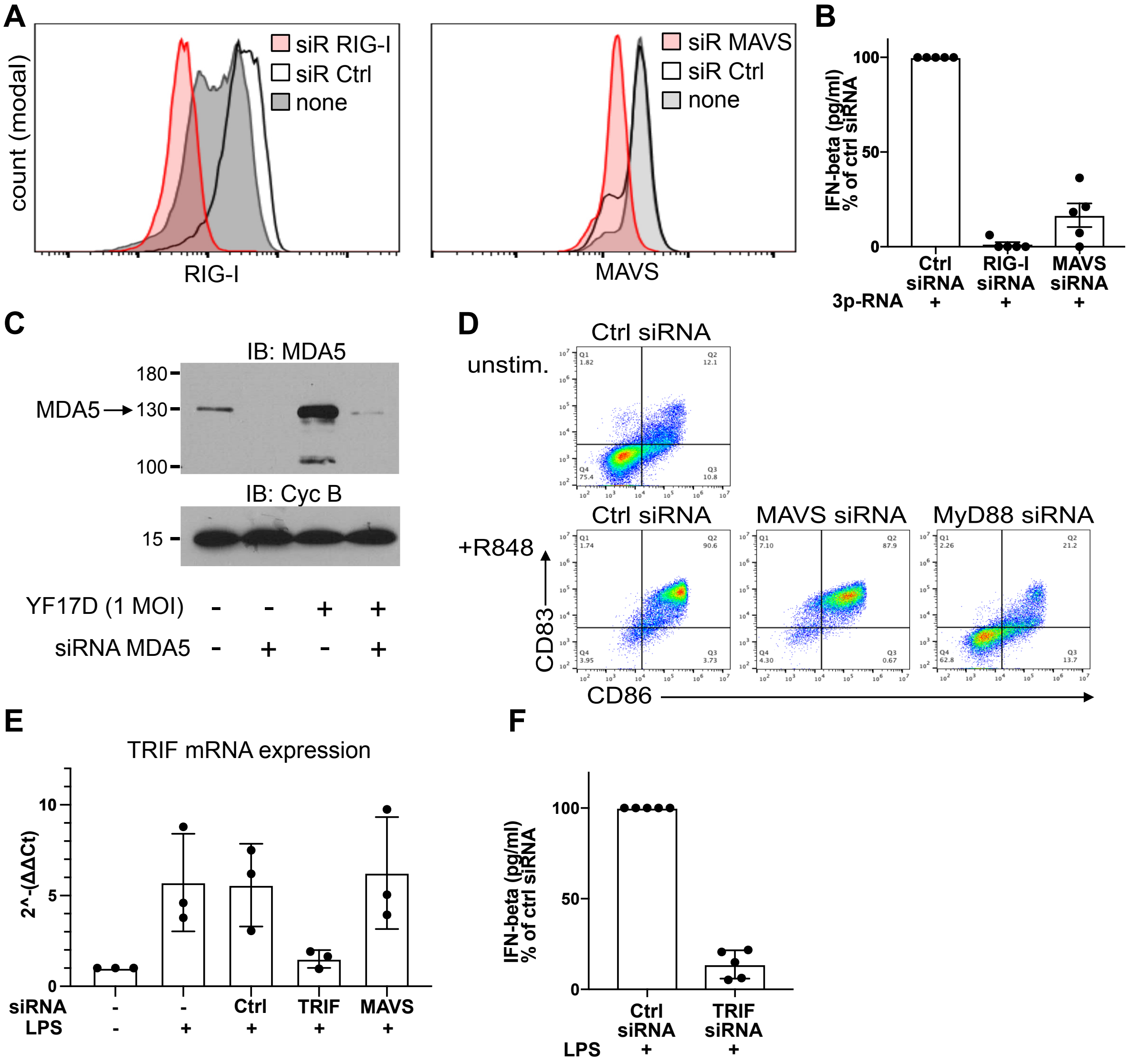


**Figure S4 Confirmation of knockdown in human mo-DCs**

(A) Mo-DCs were electroporated with siRNA against RIG-I or MAVS or with control siRNA as indicated. After 48 hrs mo-DCs were infected with YF17D at 1 MOI and cultured for additional 36 hrs. Cells were then stained intracellularly for RIG-I and MAVS and analyzed by flow cytometry. Representative results are shown as overlay histgrams (grey: untreated, white: control siRNA, red: RIG-I or MAVS-siRNA). (B) IFN-β concentrations were measured in the supernatants of mo-DCs after treatment with the indicated siRNAs and stimulation with 5 µg/ml 3pRNA (5'ppp-dsRNA from Invivogen) transfected with lipofectamine 2000 (Thermo Fisher Scientific). Results are shown as percentage of the control siRNA condition (mean +/- SD, n=5). (C) mo-DCs were transfected with siRNA against MDA5 or untreated and 48 hrs later were infected with YF17D at 1 MOI for 36 hrs or not infected. Cell lysates were separated by SDS-PAGE and MDA5 and Cyclophilin B were detected by western blot. (D)

MoDCs were transfected with control siRNA or siRNA against MAVS or MyD88 and after 48 hrs were stimulated with 3 μM R848 for 36 hrs. Mo-DCs transfected with control siRNA and cultured in medium were used as an unstimulated control. CD86 and CD83 expression were detected by flow cytometry. Representative results are shown as pseudocolor dot plots. (E) MoDCs were transfected with control siRNA, or siRNA targeting TRIF or MAVS or untreated. SMARTPool siRNA (Dharmakon) against TRIF (NM_182919) and MAVS (L-024237-00-0005) were used, as well as a control non-targeting siRNA pool (D-001810-10-05). Mo-DCs were stimulated with LPS (10 ng/ml) as indicated in the graph and analysed for relative expression of TRIF mRNA by qPCR after 48 hrs. The fold change compared to the untreated control cells is shown on the y-axis. (F) IFN-β concentrations were measured in the supernatants of mo-DCs after treatment with the indicated siRNAs and stimulation with 10 ng/ml LPS.

### Supplementary Table S1 - List of antibodies used for flow cytometry

**Antibodies used for flow cytometry Figure 1**

**PBMC panel - Figure 1A**

| Target | Clone name | Company | Catalog no. | Conjugate | Dilution |
|---|---|---|---|---|---|
| Life/Dead | | BioLegend | 423112 | Zombie Green | 1:1000 |
| CD14 | M5E2 | BioLegend | 301814 | PE-Cy7 | 1:200 |
| HLA-DR | L243 | BioLegend | 307646 | BV510 | 1:50 |
| CD123 | 6H6 | eBioscience | 45-1239-42 | PerCP-Cy5.5 | 1:100 |
| CD3 | OKT3 | eBioscience | 17-0048-42 | APC | 1:100 |
| CD19 | HIB19 | BioLegend | 302237 | BV650 | 1:200 |
| CD16 | 3G8 | BioLegend | 302026 | AlexaFluor 700 | 1:100 |
| CD56 | 5.1H11 | BioLegend | 362552 | BV421 | 1:40 |
| CD88 | S5/1 | BioLegend | 344303 | PE | 1:50 |
| CD11c | BU15 | BioLegend | 337228 | PE Dazzle594 | 1:100 |
| CD86 | IT2.2 | BioLegend | 305430 | BV605 | 1:100 |
| YF17D-4G2 | D1-4G2-4-15 | NOVUS | NBP2-52709AF647 | AF647 | 1:50 |

**mo-DC YF17D-Venus panel Figure 1B**

| Target | Clone | Company | Catalog no. | Conjugate | Dilution |
|---|---|---|---|---|---|
| Viability | | BioLegend | 423106 | Zombie NIR | 1:1000 |
| CD14 | M5E2 | BioLegend | 301814 | Pe-Cy7 | 1:200 |
| CD1a | HI149 | eBioscience | 14-0019-82 | PE | 1:100 |
| CD86 | IT2.2 | BioLegend | 305430 | BV605 | 1:200 |

**moDC YF17D panel - Figure 1B**

| Target | Clone | Company | Catalog no. | Conjugate | Dilution |
|---|---|---|---|---|---|
| YF17D-4G2 | D1-4G2-4-15 | NOVUS | NBP2-52709AF647 | AlexaFluor 64 | 1:50 |
| Viability | | BioLegend | 423106 | Zombie NIR | 1:1000 |
| CD14 | M5E2 | BioLegend | 301814 | Pe-Cy7 | 1:200 |
| CD1a | HI149 | eBioscience | 14-0019-82 | PE | 1:100 |
| CD86 | IT2.2 | BioLegend | 305430 | BV605 | 1:200 |

**Antibodies used for flow cytometry Figure 2A, B, E, F**

| Target | Clone name | Company | Catalogue no. | Fluorophore | Dilution | Comment |
|---|---|---|---|---|---|---|
| CD1a | HI149 | eBioscience | 48-0019-42 | eF450 | 1:200 | |
| HLA-DR | L243 | BioLegend | 307646 | BV510 | 1:50 | |
| CD86 | GL-1 | BioLegend | 105036 | BV605 | 1:100 | |
| CD80 | L307.4 | BD Biosciences | 564158 | BV650 | 1:50 | |
| PD-L1 | MIH1 | eBioscience | 15-5983-42 | PE Cy5 | 1:100 | |
| CD88 | S5/1 | BioLegend | 344318 | PE dazzle 594 | 1:50 | monocytes |
| CD11c | Bu15 | BioLegend | 337228 | PE dazzle 594 | 1:200 | mo-DCs |
| CD14 | M5E2 | BioLegend | 301814 | PE Cy7 | 1:200 | |
| CD16 | 3G8 | BioLegend | 302026 | AF700 | 1:100 | |
| CD83 | HB15e | BioLegend | 305312 | APC | 1:50 | |
| Live | | BioLegend | 423106 | Zombie NIR | 1:1000 | |

**Antibodies used for flow cytometry Fig. 2C,D**

| Target | Clone | Company | Catalog no. | Conjugate | Dilution |
|---|---|---|---|---|---|
| Viability | | BioLegend | 423106 | Zombie NIR | 1:1000 |
| CD14 | M5E2 | BioLegend | 301814 | Pe-Cy7 | 1:200 |
| CD1a | HI149 | eBioscience | 14-0019-82 | PE | 1:100 |
| CD86 | IT2.2 | BioLegend | 305430 | BV605 | 1:200 |

**Antibodies used for flow cytometry Figure 3**

| Target | Clone name | Company | Catalogue no. | Fluorophore | Dilution |
|---|---|---|---|---|---|
| CD3 | SK7 | BD Biosciences | 340948 | PerCP-Cy5.5 | 1 to 200 |
| CD4 | OKT4 | BioLegend | 317426 | AF700 | 1 to 320 |
| CD8 | SK1 | BioLegend | 344730 | BV650 | 1 to 320 |
| CD11c | 3.9 | BioLegend | 301608 | PE Cy7 | 1 to 100 |
| CD14 | M5E2 | BioLegend | 301834 | BV605 | 1 to 100 |
| CD40L | 24-31 | BioLegend | 310806 | PE | 1 to 50 |
| CD86 | IT2.2 | BioLegend | 305440 | BV711 | 1 to 100 |
| HLADR | L243 | BioLegend | 307654 | PE CF594 | 1 to 100 |
| IFNg | B27 | BD Biosciences | 560372 | V450 | 1 to 80 |
| TNFa | MAb11 | BioLegend | 502912 | APC | 1 to 80 |

**Antibodies used for flow cytometry Figure 4**

| Target | Clone name | Company | Catalogue no. | Fluorophore | Dilution |
|---|---|---|---|---|---|
| CD11b | M1/70 | BioLegend | 101243 | BV785 | 1:200 |
| CD11c | N418 | BioLegend | 117307 | PB | 1:200 |
| F4/80 | BM8 | BioLegend | 123115 | APC | 1:200 |
| MHC-II | AF6-120.01 | BioLegend | 116407 | PE | 1:200 |

### Supplementary Methods

#### Detection of RIG-I and MAVS by intracellular staining and flow cytometry

Mo-DCs were fixed using BD Cytofix fixation buffer (BD Biosciences, USA) for 15 min at room temperature and washed once in permeabilization buffer (PBS, 0.5% Saponin, 1% BSA, 0.01% Sodium azide). Incubation with mouse-anti-RIG-I (Alme-1, 1:1000) and anti-MAVS-AF647 (E-3, 1:200) respectively was performed in permabilization buffer for 20 minutes at RT. For RIG-I staining cells were washed in permabilization buffer and incubated with goat-anti-mouse IgG-PE (SouthernBiotech, 1:500) for 20 min at RT and washed again in permeabilization buffer followed by washing in PBS.  The samples were measured on a CytoFLEX S flow cytometer (Beckman Coulter) and data was analysed using FlowJo V10.5.3 (FlowJo LLC, USA).

#### MDA5 immunoblot

Whole cell lysates were prepared and separated by SDS-PAGE on a 10% acrylamide gel. Proteins were transferred onto nitrocellulose membrane. After blocking with TBST/5% milk powder the membrane was incubated with anti-MDA5 antibody (Abcam ab126630 1:1000) rinsed in TBST buffer and incubated with HRP-coupled goat anti rabbit secondary antibody (Jackson Immuno Research 1:7500). HRP activity was detected by Dura chemiluminescence reagent (Thermo Fisher Scientific).

#### Relative quantification of TRIF mRNA expression by qPCR

RNA was isolated from mo-DCs using Trizol reagent (Thermo Fisher Scientific) with 0.5 mL/sample following the manufacturers protocol and incubated with RNAase-free DNAase I (Thermofisher Scientific) to remove genomic DNA. Purified RNA was then reverse transcribed using the Superscript III reverse transcriptase (ThermoFisher Scientific). cDNA was diluted 1:5 in nuclease-free water, and qPCR was performed with the SYBR Green method on a Roche Light Cycler 480. The following primers were used: hTRIF (forward) GTCATCTGCCACTTTCAGGA, hTRIF (reverse) CACCGTATCCAGTTCTGACC, GAPDH (forward) ACATCGCTCAGACACCATG, GAPDH (reverse) TGTAGTTGAGGTCAATGAAGGG. Data were normalized relative to the expression of the reference gene GAPDH. The 2^-△△CT method was used for relative mRNA expression analysis and to calculate the fold-changes by normalizing to the medium control.
